# Local PCA Shows How the Effect of Population Structure Differs Along the Genome

**DOI:** 10.1101/070615

**Authors:** Han Li, Peter Ralph

## Abstract

Population structure leads to systematic patterns in measures of mean relatedness between individuals in large genomic datasets, which are often discovered and visualized using dimension reduction techniques such as principal component analysis (PCA). Mean relatedness is an average of the relationships across locus-specific genealogical trees, which can be strongly affected on intermediate genomic scales by linked selection and other factors. We show how to use local principal components analysis to describe this meso-scale heterogeneity in patterns of relatedness, and apply the method to genomic data from three species, finding in each that the effect of population structure can vary substantially across only a few megabases. In a global human dataset, localized heterogeneity is likely explained by polymorphic chromosomal inversions. In a range-wide dataset of *Medicago truncatula*, factors that produce heterogeneity are shared between chromosomes, correlate with local gene density, and may be caused by linked selection, such as background selection or local adaptation. In a dataset of primarily African *Drosophila melanogaster*, large-scale heterogeneity across each chromosome arm is explained by known chromosomal inversions thought to be under recent selection, and after removing samples carrying inversions, remaining heterogeneity is correlated with recombination rate and gene density, again suggesting a role for linked selection. The visualization method provides a flexible new way to discover biological drivers of genetic variation, and its application to data highlights the strong effects that linked selection and chromosomal inversions can have on observed patterns of genetic variation.

## 1 Introduction

Wright [68] defined *population structure* to encompass “such matters as numbers, composition by age and sex, and state of subdivision”, where “subdivision” refers to restricted migration between subpopulations. The phrase is also commonly used to refer to the genetic patterns that result from this process, as for instance reduced mean relatedness between individuals from distinct populations. However, it is not necessarily clear what aspects of demography should be included in the concept. For instance, Blair [4] defines *population structure* to be the sum total of “such factors as size of breeding populations, periodic fluctuation of population size, sex ratio, activity range and *differential survival of progeny*” (emphasis added). The definition is similar to Wright’s, but differs in including the effects of natural selection. On closer examination, incorporating differential survival or fecundity makes the concept less clear: should a randomly mating population consisting of two types that are partially reproductively isolated from each other be said to show population structure or not? Whatever the definition, it is clear that due to natural selection, the effects of population structure – the *realized* patterns of genetic relatedness – differ depending on which portion of the genome is being considered. For instance, strongly locally adapted alleles of a gene will be selected against in migrants to different habitats, increasing genetic differentiation between populations near to this gene. Similarly, newly adaptive alleles spread first in local populations. These observations motivate many methods to search for genetic loci under selection, as for example in Huerta-Sánchez et al. [30], Martin et al. [46], and Duforet-Frebourg et al. [19].

These realized patterns of genetic relatedness summarize the shapes of the genealogical trees at each location along the genome. Since these trees vary along the genome, so does relatedness, but averaging over sufficiently many trees we hope to get a stable estimate that doesn’t depend much on the genetic markers chosen. This is not guaranteed: for instance, relatedness on sex chromosomes is expected to differ from the autosomes; and positive or negative selection on particular loci can dramatically disort shapes of nearby genealogies [3, 10, 36]. Indeed, many species show chromosome-scale variation in diversity and divergence (e.g., Langley et al. [40]); species phylogenies can differ along the genome due to incomplete lineage sorting, adaptive introgression and/or local adaptation (e.g., Ellegren et al. [21], Nadeau et al. [49], Pease and Hahn [55], Pool [57], Vernot and Akey [65]); and theoretical expectations predict that geographic patterns of relatedness should depend on selection [14].

Patterns in genome-wide relatedness are often summarized by applying principal components analysis (PCA, Patterson et al. [54]) to the genetic covariance matrix, as pioneered by Menozzi et al. [48]. The results of PCA can be related to the genealogical history of the samples, such as time to most recent common ancestor and migration rate between populations [47, 51], and sometimes produce “maps” of population structure that reflect the samples’ geographic origin distorted by rates of gene flow [52].

Modeling such “background” kinship between samples is essential to genome-wide association studies (GWAS, Astle and Balding [2], Price et al. [59]), and so understanding variation in kinship along the genome could lead to more generally powerful methods, and may be essential for doing GWAS in species with substantial heterogeneity in realized patterns of mean relatedness along the genome.

Others have applied PCA to windows of the genome: Ma and Amos [43] used local PCA much as we do to identify putative chromosomal inversions. Bryc et al. [7] and Brisbin et al. [6] used PCA to infer tracts of local ancestry in recently admixed populations, but by projecting each genomic window onto the axes of a single, globally-defined PCA rather than doing PCA separately on each window.

A note on nomenclature: In this work we describe variation in patterns of relatedness using local PCA, where “local” refers to proximity along the genome. A number of general methods for dimensionality reduction also use a strategy of “local PCA” (e.g., Kambhatla and Leen [34], Manjón et al. [45], Roweis and Saul [60], Weingessel and Hornik [67]), performing PCA not on the entire dataset but instead on subsets of observations, providing local pictures which are then stitched back together to give a global picture. At first sight, this differs from our method in that we restrict to subsets of *variables* instead of subsets of observations. However, if we flip perspectives and think of each genetic variant as an observation, our method shares common threads, although our method does not subsequently use adjacency along the genome, as we aim to identify similar regions that may be distant.

It is common to describe variation along the genome of simple statistics such as *F*_*ST*_ and to interpret the results in terms of the action of selection (e.g., Ellegren et al. [21], Turner et al. [64]). However, a given pattern (e.g., valleys of *F*_*ST*_) can be caused by more than one biological process [8, 18], which in retrospect is unsurprising given that we are using a single statistic to describe a complex process. It is also common to use methods such as PCA to visualize large-scale patterns in mean genome-wide relatedness. In this paper we show if and how patterns of mean relatedness vary systematically along the genome, in a way particularly suited to large samples from geographically distributed populations. Geographic population structure sets the stage by establishing “background” patterns of relatedness; our method then describes how this structure is affected by selection and other factors. Our aim is not to identify outlier loci, but rather to describe larger-scale variation shared by many parts of the genome; correlation of this variation with known genomic features can then be used to uncover its source.

## 2 Materials and Methods

As depicted in Figure 1, the general steps to the method are: (1) divide the genome into windows, (2) summarize the patterns of relatedness in each window, (3) measure dissimilarity in relatedness between each pair of windows, (4) visualize the resulting dissimilarity matrix using multidimensional scaling (MDS), and (5) combine similar windows to more accurately visualize local effects of population structure using PCA.

**Figure 1:**
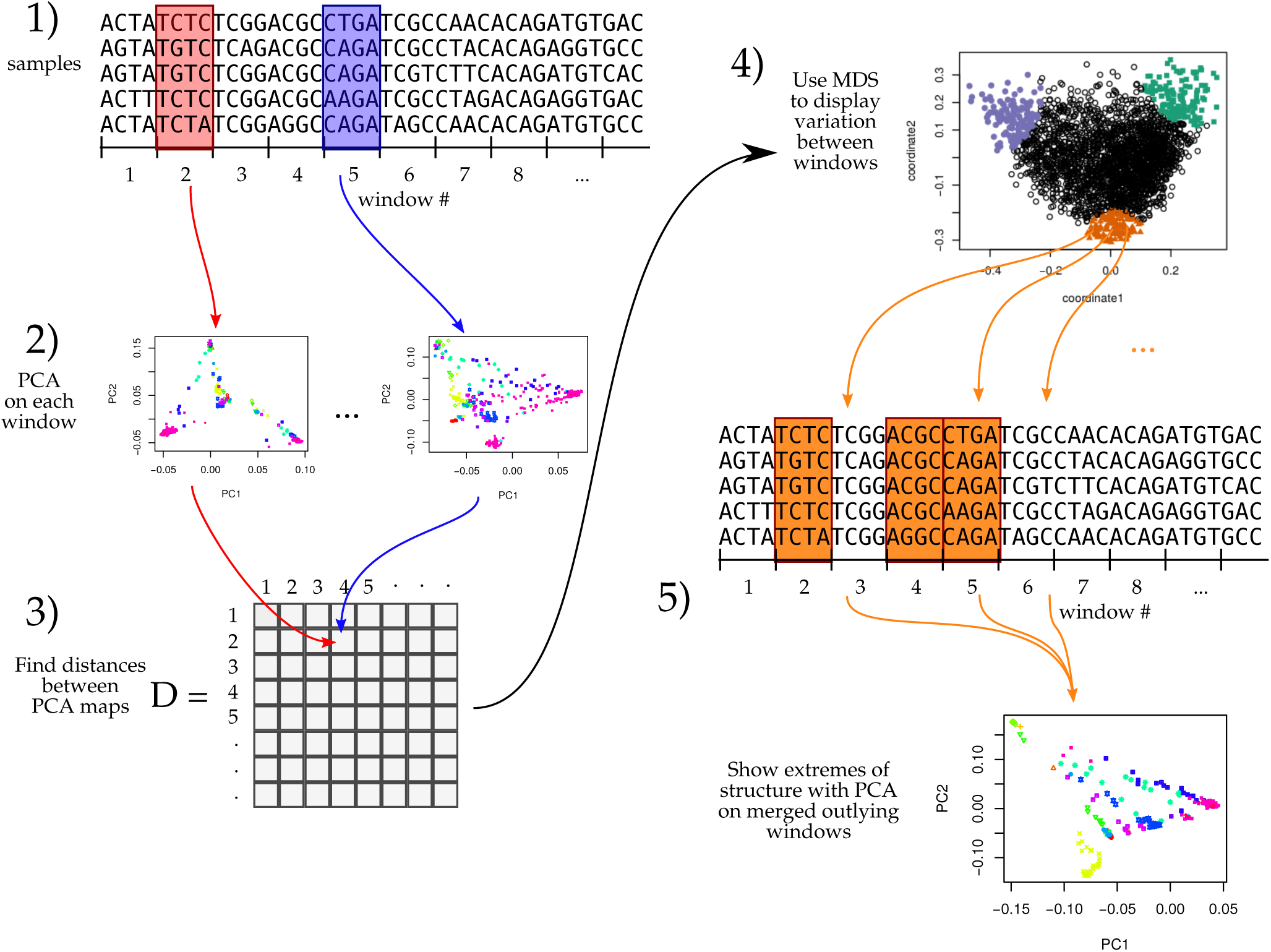
An illustration of the method; see Methods for details.

### 2.1 PCA in genomic windows

To begin, we first recoded sampled genotypes as numeric matrices in the usual manner, by recording the number of nonreference alleles seen at each locus for each sample. We then divided the genome into contiguous segments (“windows”) and applied principal component analysis (PCA) as described in McVean [47] separately to the submatrices that corresponded to each window. The choice of window length entails a tradeoff between signal and noise, since shorter windows allow better resolution along the genome but provide less precise estimates of relatedness. A method for choosing a window length to balance these considerations is given in Appendix A. Precisely, denote by *Z* the *L* × *N* recoded genotype matrix for a given window (*L* is the number of SNPs and *N* is the sample size), and by 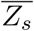 the mean of non-missing entries for allele *s*, so that 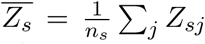, where the sum is over the *n*_*s*_ nonmissing genotypes. We first compute the mean-centered matrix *X*, as 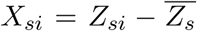, and preserving missingness. (This mean-centering makes the result not depend on the choice of reference allele, exactly if there is no missing data, and approximately otherwise.) Next, we find the covariance matrix of *X*, denoted *C*, as 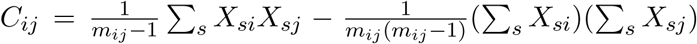, where all sums are over the *m*_*ij*_ sites where both sample *i* and sample *j* have nonmissing genotypes. The principal components are the eigenvectors of *C*, normalized to have Euclidean length equal to one, and ordered by magnitude of the eigenvalues.

The top 2–5 principal components are generally good summaries of population structure; for ease of visualization we usually only use the first two (referred to as *PC*1 and *PC*2), and check that results hold using more. The above procedure can be performed on any subset of the data; for future reference, denote by *PC*1_*j*_ and *PC*2_*j*_ the result after applying to all SNPs in the *j*^th^ window. (Note, however, that our measure of dissimilarity between windows does not depend on PC ordering.)

### 2.2 Similarity of patterns of relatedness between windows

We think of the local effects of population structure as being summarized by *relative* position of the samples in the space defined by the top principal components. However, we do not compare patterns of relatedness of different genomic regions by directly comparing the PCs, since rotations or reflections of these imply identical patterns of relatedness. Instead, we compare the low-dimensional approximations of the local covariance matrices obtained using the top *k* PCs, which is invariant under reordering of the PCs, reflections, and rotations and yet contains all other information about the PCs. (For results shown here, we use *k* = 2.) Furthermore, to remove the effect of artifacts such as mutation rate variation, we also rescale each approximate covariance matrix to be of similar size (precisely, so that the underlying data matrix has trace norm equal to one).

To do this, define the *N* × *k* matrix *V* (*i*) so that *V* (*i*)_*· ℓ*_, the *ℓ*^th^ column of *V* (*i*), is equal to the *ℓ*^th^ principal component of the *i*^th^ window, multiplied by 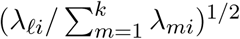, where λ_*ℓi*_ is the *ℓ*^th^ eigenvalue of the genetic covariance matrix. Then, the rescaled, rank *k* approximate covariance matrix for the *i*^th^ window is

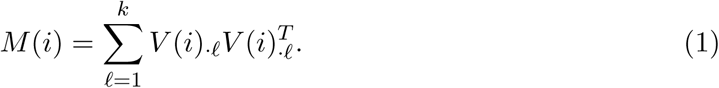

To measure the similarity of patterns of relatedness for the *i*^th^ window and *j*^th^ window, we then use Euclidean distance *D*_*ij*_ between the matrices *M* (*i*) and *M* (*j*), defined by 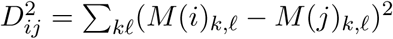.

The goal of comparing PC plots up to rotation and reflection turned out to be equivalent to comparing rank-*k* approximations to local covariance matrices. This suggests instead directly comparing entire local covariance matrices. However, with thousands of samples and tens of thousands of windows, computing the distance matrix would take months of CPU time, while as defined above, *D* can be computed in minutes using the following method. Since for square matrices *A* and *B*, 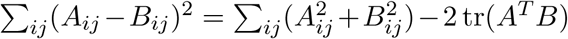, then due to the orthogonality of eigenvectors and the cyclic invariance of trace, *D*_*ij*_ can be computed efficiently as

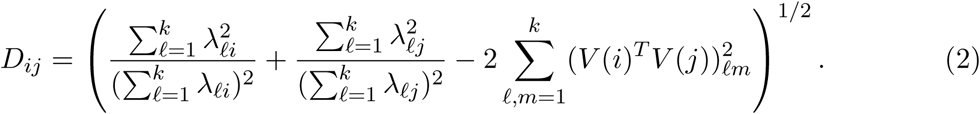

### 2.3 Visualization of results

We use multidimensional scaling (MDS) to visualize relationships between windows as summarized by the dissimilarity matrix *D*. MDS produces a set of *m* coordinates for each window that give the arrangement in *m*-dimensional space that best recapitulates the original distance matrix. For results here, we use *m* = 2 to produce one-or two-dimensional visualizations of relationships between windows’ patterns of relatedness.

We then locate variation in patterns of relatedness along the genome by choosing collections of windows that are nearby in MDS coordinates, and map their positions along the genome. A visualization of the effects of population structure across the entire collection is formed by extracting the corresponding genomic regions and performing PCA on all, aggregated, regions.

### 2.4 Testing

We tested the method using two types of simulation. First, to verify expected behavior, we simulated “genomes” as an independent sequence of correlated Gaussian “genotypes”, using a different covariance matrix in the first quarter, middle half, and last quarter of the chromosome. The details of the simulation, also designed to detect sensitivity to PC switching, are given in Appendix B.1. To verify robustness to missing data, we ran the method after randomly dropping 50% of the genotypes in the first half of the genome; if the method is misled by missing data, then it will distinguish the two halves of the chromosome rather than the segments having different covariance matrices.

To provide a realistic test, we next used forwards-time, individual-based simulations, implemented using SLiM v3 [25], which are described in detail in Appendix B.2. To provide realistic population structure for PCA to identify, each simulation had at least 5,000 diploid individuals, living across a continuous square range, with Gaussian dispersal and local density-dependent competition. Each genome was modeled on human chromosome 7, which is 1.54 × 10^8^bp long, with an overall recombination rate of 1.6785 crossovers per chromosome per generation. To improve speed, we used tskit [35] to record tree sequences in SLiM [26] and to add neutral mutations afterwards, at a rate of 10^−9^ per bp per generation. Most simulations were neutral, but we also included linked selection, of two types. First, we introduced selected mutations into two regions, which extended from 1/3 to 1/2 and from 5/6 to the end of the genome respectively. These had selection coefficients from a Gamma distribution with shape 2 and mean 0.005 at a rate of 10^−10^ per bp, that were either beneficial (with probability 1/30) or deleterious (otherwise). Second, to roughly model a recent expansion followed by local adaptation, we introduced mutations in the same manner as above, except that mutations were no longer unconditionally deleterious or beneficial: each selection coefficient was multiplied by a factor depending on the spatial location of the individual being evaluated, varying linearly from -1 at the left side of the range to +1 at the right edge. In all simulations, genome-wide PCA displayed a map of the population range, as expected.

### 2.5 Datasets

We applied the method to genomic datasets with good geographic sampling: 380 African *Drosophila melanogaster* from the Drosophila Genome Nexus [39], a worldwide dataset of humans, 3,965 humans from several locations worldwide from the POPRES dataset [50], and 263 *Medicago truncatula* from 24 countries around the Mediterranean basin a rangewide dataset of the partially selfing weedy annual plant from the *Medicago truncatula* Hapmap Project [63], as summarized in Table 1.

**Table 1:** Descriptive statistics for each dataset used.

| species | # SNPs per window | mean window length (bp) | mean # windows per chromosome | mean % variance explained by top 2 PCs |
| --- | --- | --- | --- | --- |
| *Drosophila melanogaster* | 1,000 | 9,019 | 2,674 | 0.53 |
| Human | 100 | 636,494 | 203 | 0.55 |
| *Medicago truncatula* | 10,000 | 102,580 | 467 | 0.50 |

#### Drosophila melanogaster

We used whole-genome sequencing data from the Drosophila Genome Nexus (http://www.johnpool.net/genomes.html, [39]), consisting of the Drosophila Population Genomics Project phases 1–3 [40, 58], and additional African genomes [39]. After removing 20 genomes with more than 8% missing data, we were left with 380 samples from 16 countries across Africa and Europe. Since the *Drosophila* samples are from inbred lines or haploid embryos, we treat the samples as haploid when recoding; regions with residual heterozygosity were marked as missing in the original dataset; we also removed positions with more than 20% missing data. Each chromosome arm we investigated (X, 2L, 2R, 3L, and 3R) has 2–3 million SNPs; PCA plots for each arm are shown in Figure S1.

#### Human

We also used genomic data from the entire POPRES dataset [50], which has array-derived genotype information for 447,267 SNPs across the 22 autosomes of 3,965 samples in total: 346 African-Americans, 73 Asians, 3,187 Europeans and 359 Indian Asians. Since these data derive from genotyping arrays, the SNP density is much lower than the other datasets, which are each derived from whole genome sequencing. We excluded the sex chromosomes and the mitochondria. PCA plots for each chromosome, separately, are shown in Figure S2.

#### Medicago truncatula

Finally, we used whole-genome sequencing data from the *Medicago truncatula* Hapmap Project [63], which has 263 samples from 24 countries, primarily distributed around the Mediterranean basin. Each of the 8 chromosomes has 3–5 million SNPs; PCA plots for these are shown in Figure S3. We did not use the mitochondria or chloroplasts.

### 2.6 Data access

The methods described here are implemented in an open-source R package available at https://github.com/petrelharp/local_pca, as well as scripts to perform all analyses from VCF files at various parameter settings.

Datasets are available as follows: human (POPRES) at dbGaP with accession number phs000145.v4.p2, *Medicago* at the Medicago Hapmap http://www.medicagohapmap.org/, and *Drosophila* at the Drosophila Genome Nexus, http://www.johnpool.net/genomes.html.

## 3. Results

In all three datasets: a worldwide sample of humans, African *Drosophila melanogaster*, and a rangewide sample of *Medicago truncatula*, PCA plots vary along the genome in a systematic way, showing strong chromosome-scale correlations. This implies that variation is due to meaningful heterogeneity in a biological process, since noise due to randomness in choice of local genealogical trees is not expected to show long distance correlations. Below, we discuss the results and likely underlying causes.

### 3.1 Validation

Simple non-population-based simulations with Gaussian “genotypes” showed that the method performs as expected, clearly separating regions of the genome with different underlying covariance matrices without being affected by extreme differences in amount of missing data (Supplemental Figure S4). This simulation also verifies insensitivity to ordering of top PCs, since it was performed using a covariance matrix with the top two eigenvalues equal, so that the order of empirical eigenvectors (PCs) switches randomly.

Individual-based simulations using SLiM [25] allowed us to test the effects of recombination and mutation rate variation, as well as linked selection. As expected, varying recombination rate stepwise by a factor of 64 did not induce patterns in the MDS visualizations correlated with recombination rate (Supplemental Figure S5). Since varying mutation rate with a fixed recombination map is equivalent to varying the recombination map and remapping windows, this also indicates that the method is not misled by variation in mutation rate. On the other hand, a recombination map with hotspots (the HapMap human female map for chromosome 7 [32]) induced outliers at long regions of low recombination rate (also as expected).

Simulations with linked selection produced mixed results (Figure S6). The method strongly identified the regions under spatially varying linked selection. It also identified the regions (although less unambiguously) with constant selection and stepwise varying recombination rate, but did not clearly identify them with constant recombination rate. This difference is likely because recombination rates are overall lower in the first case, leading to a stronger effect of linked selection. These tests are not meant to be comprehensive survey of linked selection, but only to demonstrate that linked selection can produce signals similar to what we see in real data.

### 3.2 Drosophila melanogaster

We applied the method to windows of average length 9 Kbp, across chromosome arms 2L, 2R, 3L, 3R and X separately. The first column of Figure 2 is a multidimensional scaling (MDS) visualization of the matrix of dissimilarities between genomic windows: in other words, genomic windows that are closer to each other in the MDS plot show more similar patterns of relatedness. For each chromosome arm, the MDS visualization roughly resembles a triangle, sometimes with additional points. Since the relative position of each window in this plot shows the similarity between windows, this suggests that there are at least three extreme manifestations of population structure typified by windows found in the “corners” of the figure, and that other windows’ patterns of relatedness may be a mixture of those extremes. The next two columns of Figure 2 respectively depict the two MDS coordinates of each window, plotted against the window’s position along the genome, to show how the plot of the first column is laid out along the genome. The patterns did not depend on the number of PCs used (see Figure S7 for the same plot with *k* = 5 PCs), and are only weakly correlated with variation in missingness (see Figure S8).

**Figure 2:**
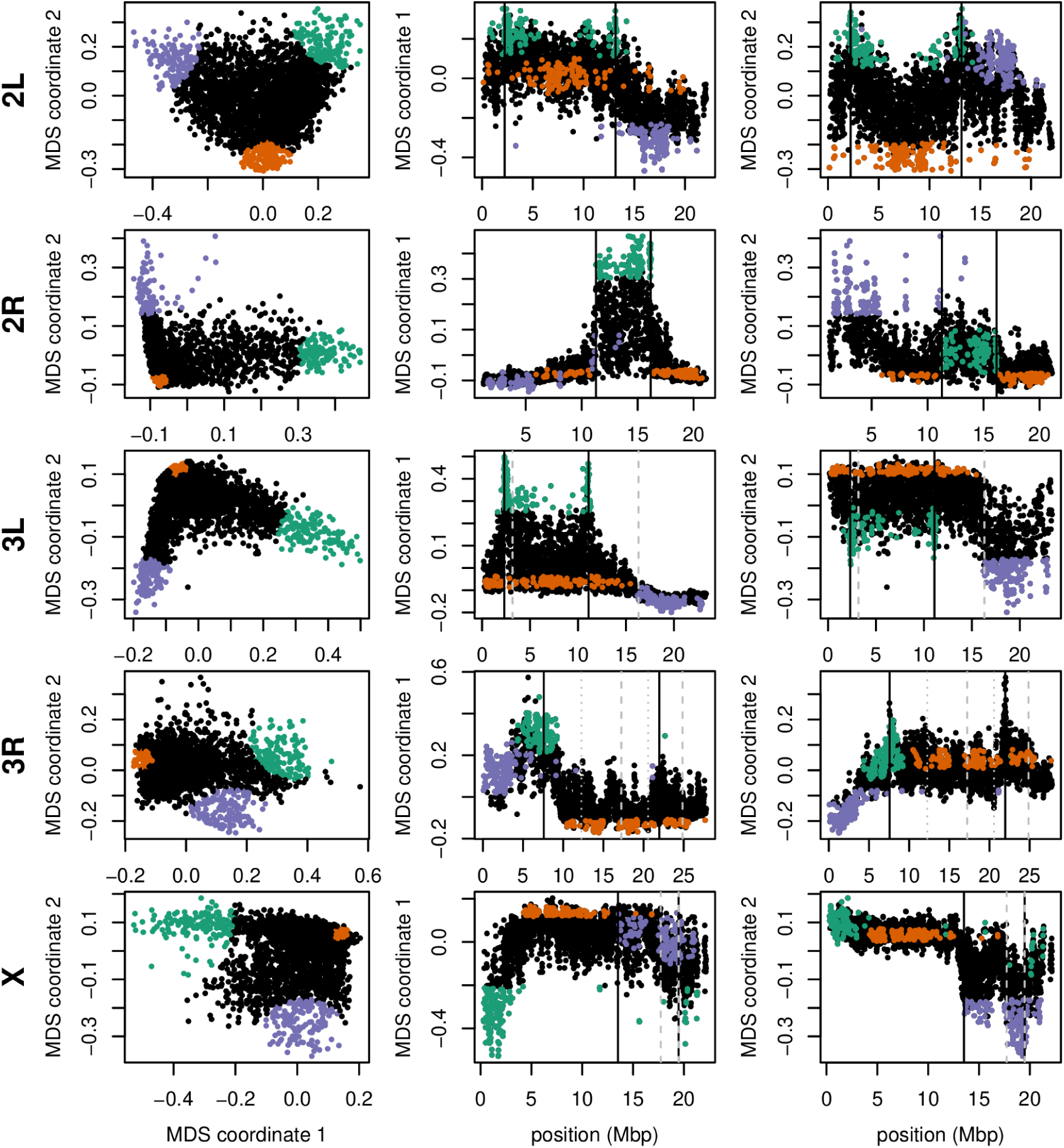
Variation in patterns of relatedness for windows across *Drosophila melanogaster* chromosome arms. In all plots, each point represents one window along the genome. The first column shows the MDS visualization of relationships between windows, and the second and third columns show the two MDS coordinates against the midpoint of each window; rows correspond to chromosome arms. Colors are consistent for plots in each row. Vertical lines show the breakpoints of known polymorphic inversions. Solid black lines are for the inversions we used in Figure 3, while dotted grey lines are for other known inversions.

To help visualize how clustered windows with similar patterns of relatedness are along each chromosome arm, we selected three “extreme” windows in the MDS plot and the 5% of windows that are closest to it in the MDS coordinates, then highlighted these windows’ positions along the genome, and created PCA plots for the windows, combined. Representative plots are shown for three groups of windows on each chromosome arm in Figure 2 (groups are shown in color), and in Supplemental Figure S9 (PCA plots). The latter plots are quite different, showing that genomic windows in different regions of the MDS plot indeed show quite different patterns of relatedness.

The most striking variation in patterns of relatedness turns out to be explained by several large inversions that are polymorphic in these samples, discussed in Corbett-Detig and Hartl [16] and Langley et al. [40]. To depict this, Figure 3 shows the PCA plots in Supplemental Figure S9 recolored by the orientation of the inversion for each sample. Taking chromosome arm 2L as an example, the two regions of similar, extreme patterns of relatedness shown in green in the first row of Figure 2 lie directly around the breakpoints of the inversion In(2L)t, and the PCA plots in the first rows of Figure 3 shows that patterns of relatedness here are mostly determined by inversion orientation. The regions shown in purple on chromosome 2L lie near the centromere, and have patterns of relatedness reflective of two axes of variation, seen in Figure 3 and Supplemental Figure S9, which correspond roughly to latitude within Africa and to degree of cosmopolitan admixture respectively (see Lack et al. [39] for more about admixture in this sample). The regions shown in orange on chromosome 2L mostly lie inside the inversion, and show patterns of relatedness that are a mixture between the other two, as expected due to recombination within the (long) inversion [24]. Similar results are found in other chromosome arms, albeit complicated by the coexistence of more than one polymorphic inversion; however, each breakpoint visibly affects patterns in the MDS coordinates (see vertical lines in Figure 2).

**Figure 3:**
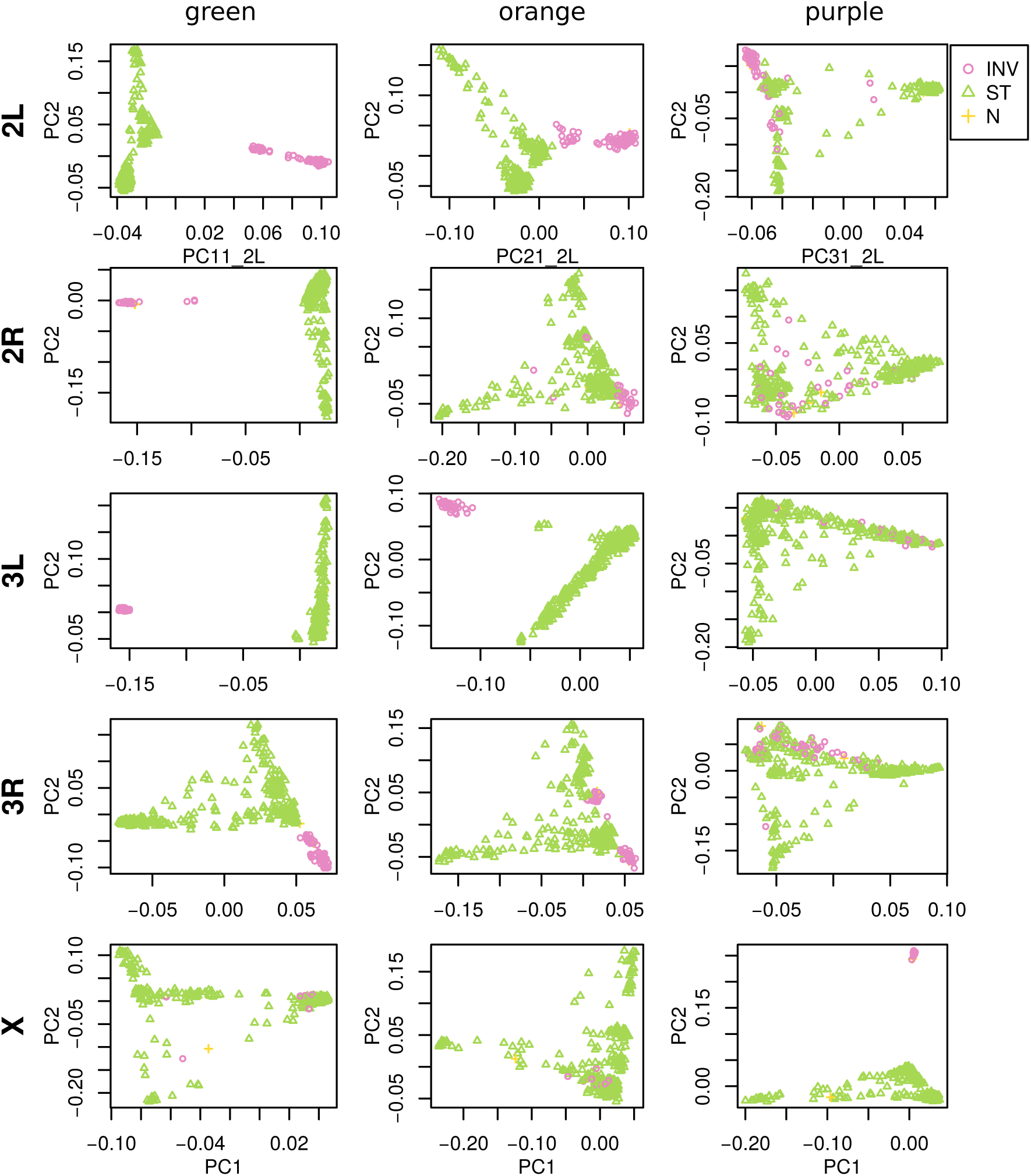
PCA plots for the three sets of genomic windows colored in Figure 2, on each chromosome arm of *Drosophila melanogaster*. In all plots, each point represents a sample. The first column shows the combined PCA plot for windows whose points are colored green in Figure 2; the second is for orange windows; and third is for purple windows. In each, samples are colored by orientation of the polymorphic inversions In(2L)t, In(2R)NS, In(3L)OK, In(3R)K and In(1)A respectively (data from [39]). In each “INV” denotes an inverted genotype, “ST” denotes the standard orientation, and “N” denotes unknown.

To see how patterns of relatedness vary in the absence of polymorphic inversions, we performed the same analyses after removing, for each chromosome arm, any samples carrying inversions on that arm. In the result, shown in Figure 4 and Supplemental Figure S10, the striking peaks associated with inversion breakpoints are gone, and previously smallerscale variation now dominates the MDS visualization. For instance, the majority of the variation along 3L in Figure 2 is on the left end of the arm, dominated by two large peaks around the inversion breakpoints; there is also a relatively small dip on the right end of the arm (near the centromere). In contrast, Supplemental Figure S10 shows that after removing polymorphic inversions, remaining structure is dominated by the dip near the centromere. Without inversions, variation in patterns of relatedness shown in the MDS plots follows similar patterns to that previously seen in *D*. *melanogaster* recombination rate and diversity [40, 44]. Indeed, correlations between the recombination rate in each window and the position on the first MDS coordinate are highly significant (Spearman’s *ρ* = 0.54, *p <* 2 × 10^−16^; Figures 4 and S11). This is consistent with the hypothesis that variation is due to selection, since the strength of linked selection increases with local gene density, measured in units of recombination distance. The number of genes – measured as the number of transcription start and end sites within each window – was not significantly correlated with MDS coordinate (*p* = 0.22).

**Figure 4:**
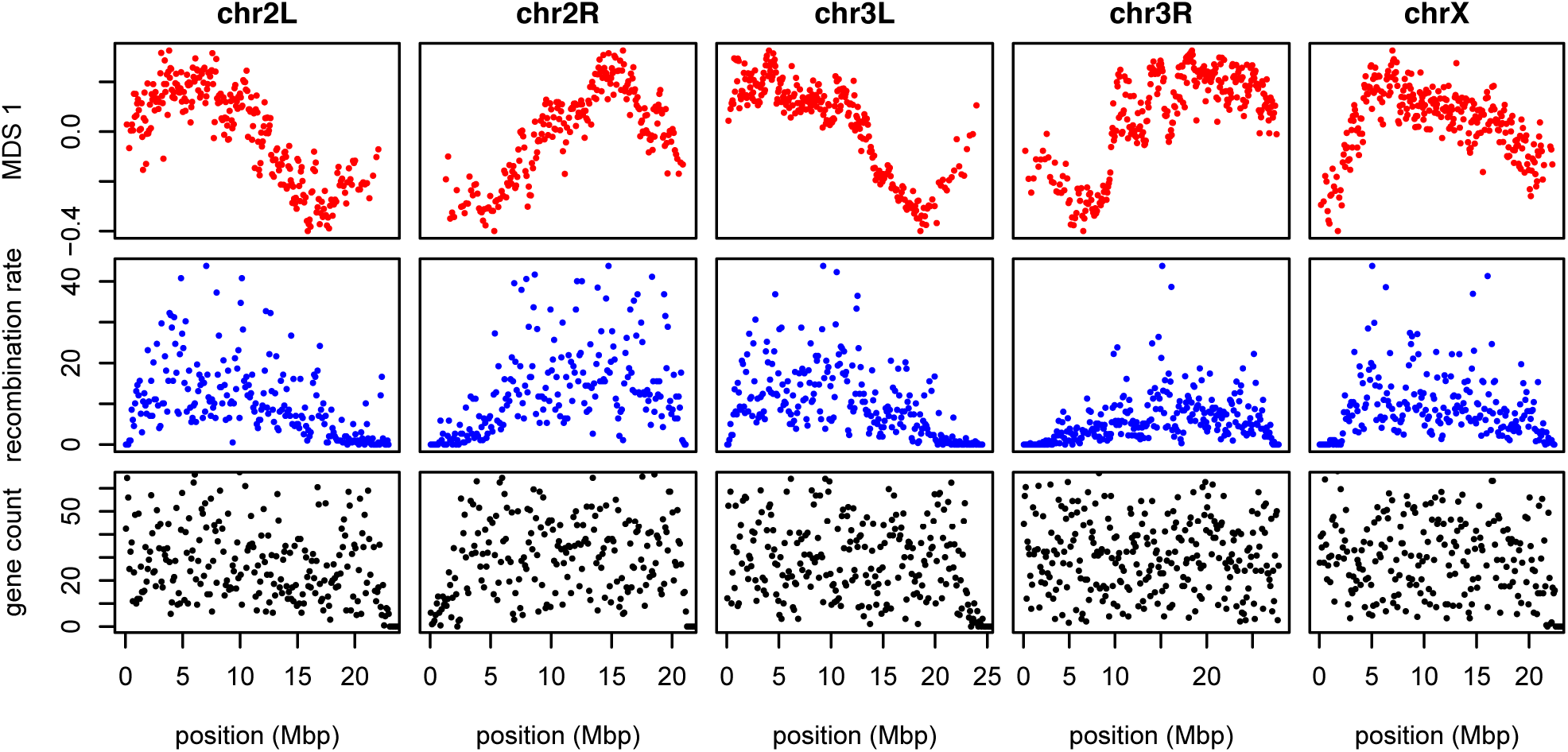
The effects of population structure without inversions is correlated to recombination rate in *Drosophila melanogaster*. The first plot (in red) shows the first MDS coordinate along the genome for windows of 10,000 SNPs, obtained after removing samples with inversions. (A plot analogous to Figure 2 is shown in Supplemental Figure S10.) The second plot (in blue) shows local average recombination rates in cM/Mbp, obtained as midpoint estimates for 100Kbp windows from the Drosophila recombination rate calculator [22] release 5, using rates from Comeron et al. [15]. The third plot (in black) shows the number of genes’ transcription start and end sites within each 100Kbp window, divided by two. Transcription start and end sites were obtained from the RefGene table from the UCSC browser. The histone gene cluster on chromosome arm 2L is excluded.

### 3.3 Human

As we did for the *Drosophila* data, we applied our method separately to all 22 human autosomes. On each, variation in patterns of relatedness was dominated by a small number of windows having similar patterns of relatedness to each other that differed dramatically from the rest of the chromosome. These may be primarily inversions: outlying windows coincide with three of the six large polymorphic inversions described in Antonacci et al. [1], notably a particularly large, polymorphic inversion on 8p23 (Figure 5). Similar plots for all chromosomes are shown in Supplementary Figures S12, S13, and S14. PCA plots of many outlying windows show a characteristic trimodal shape (shown for chromosome 8 in Figure S15), presumably distinguishing samples having each of the three diploid genotypes for each inversion orientation (although we do not have data on orientation status). This trimodal shape has been proposed as a method to identify inversions [43], but distinguishing this hypothesis from others, such as regions of low recombination rate, would require additional data.

**Figure 5:**
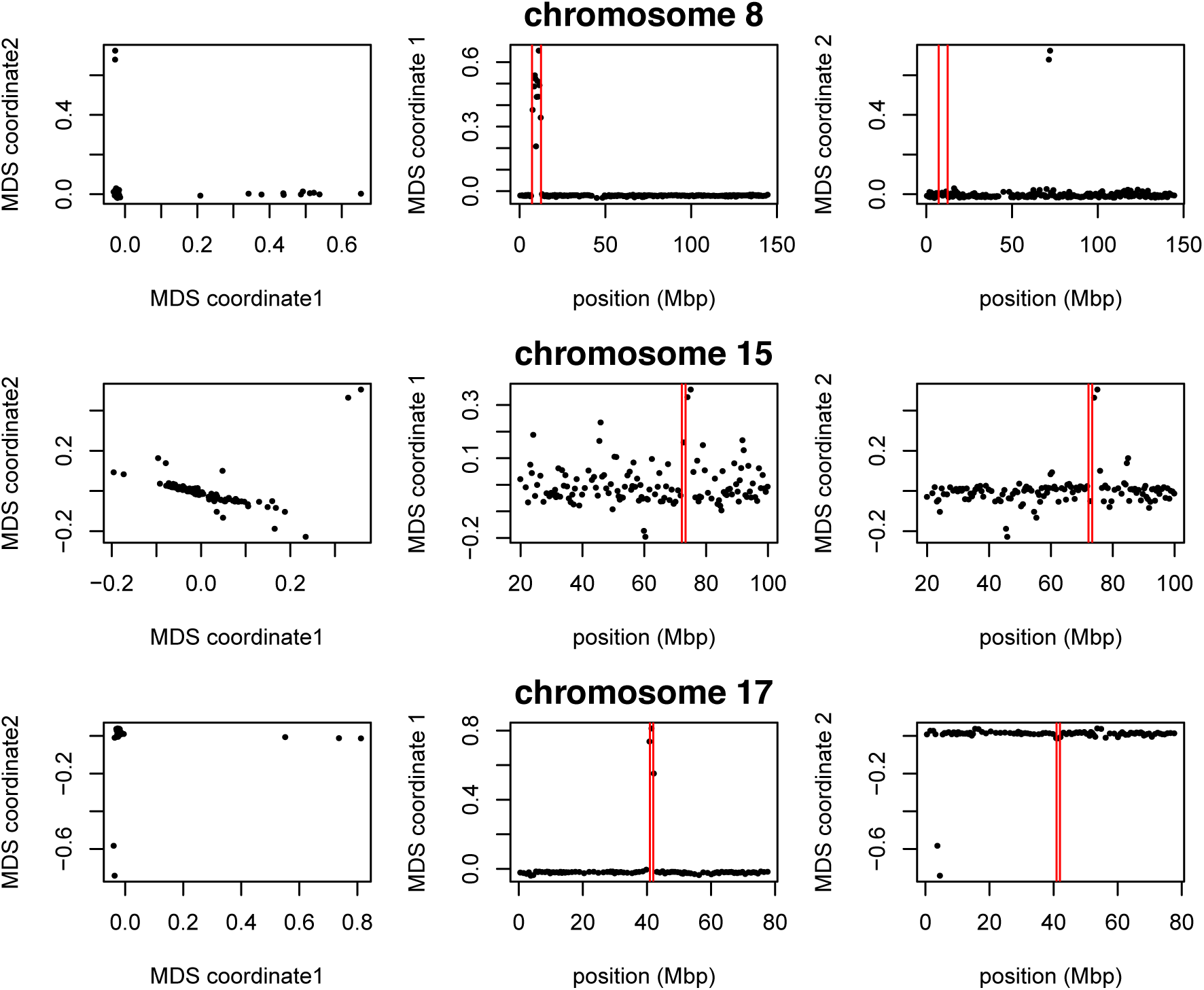
Variation in structure between windows on human chromosomes 8, 15, and 17. Each point in each plot represents a window. The first column shows the MDS visualization of relationships between windows; the second and third columns show the two MDS coordinates of each window against its position (midpoint) along the chromosome. Rows, from top to bottom show chromosomes 8, 15, and 17. The vertical red lines show the breakpoints of known inversions from Antonacci et al. [1].

We also applied the method on all 22 autosomes together, and found that, remarkably, the inversion on chromosome 8 is still the most striking outlying signal (Figure S16). Further investigation with a denser set of SNPs, allowing a finer genomic resolution, may yield other patterns.

### 3.4 Medicago truncatula

Unlike the other two species, the method applied separately on all eight chromosomes of *Medicago truncatula* showed similar patterns of gradual change in patterns of relatedness across each chromosome, with no indications of chromosome-specific patterns. This consistency suggests that the factor affecting the population structure for each chromosome is the same, as might be caused by varying strengths of linked selection. To verify that variation in the effects of population structure is shared across chromosomes, we applied the method to all chromosomes together. Results for chromosome 3 are shown in Figure 6, and other chromosomes are similar: across chromosomes, the high values of the first MDS coordinate coincide with the position of the heterochromatic regions surrounding the centromere, which often have lower gene density and may therefore be less subject to linked selection. To verify that this is a possible explanation, we counted the number of genes found in each window using gene models in Mt4.0 from jcvi.org [63], which are shown juxtaposed with the first MDS coordinate of each window in Figure 7, and are significantly correlated, as shown in Supplemental Figure S17. (Values shown are the number of start and end positions of each predicted mRNA transcript, divided by two, assigned to the nearest window.) However, other genomic features, such as distance to centromere show roughly the same patterns, so we cannot rule out alternative hypotheses. In particular, fine-scale recombination rate estimates are not available in a form mappable to Mt4.0 coordinates (although those in Paape et al. [53] appear visually similar).

**Figure 6:**
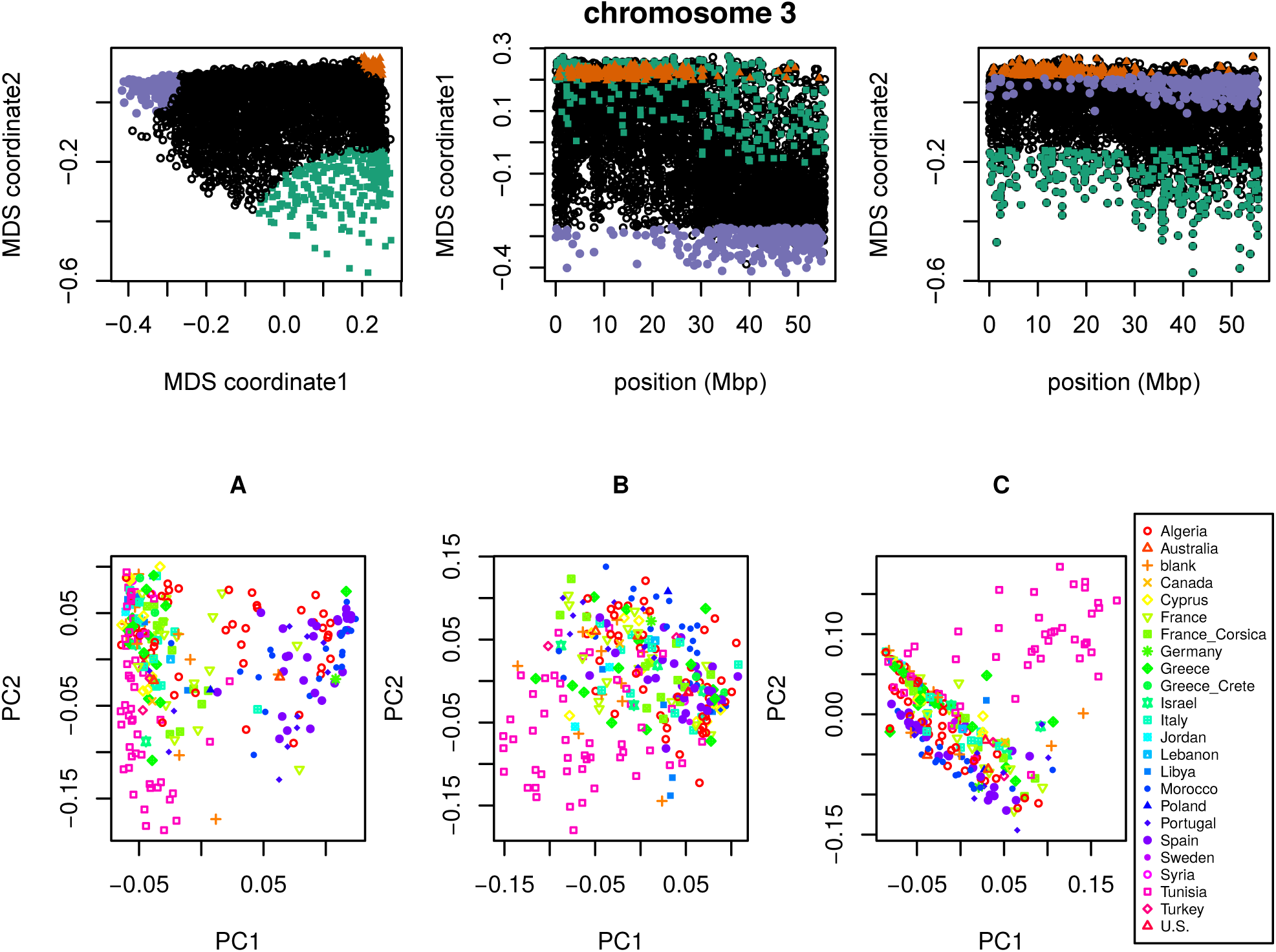
**(top)** MDS visualization of patterns of relatedness on *M*. *truncatula* chromosome 3, with corresponding PCA plots. Each point in the plot represents a window; the structure revealed by the MDS plot is strongly clustered along the chromosome, with windows in the upper-right corner of the MDS plot (colored red) clustered around the centromere, windows in the upper-left corner (purple) furthest from the centromere, and the remaining corner (green) intermediate. Plots for remaining chromosomes are shown in Supplemental Figure S18. **(bottom)** PCA plots for the sets of genomic windows colored (A) green, (B) orange, and (C) purple in the top figure. Each point corresponds to a sample, colored by country of origin. Plots for remaining chromosomes are shown in Supplemental Figure S19.

**Figure 7:**
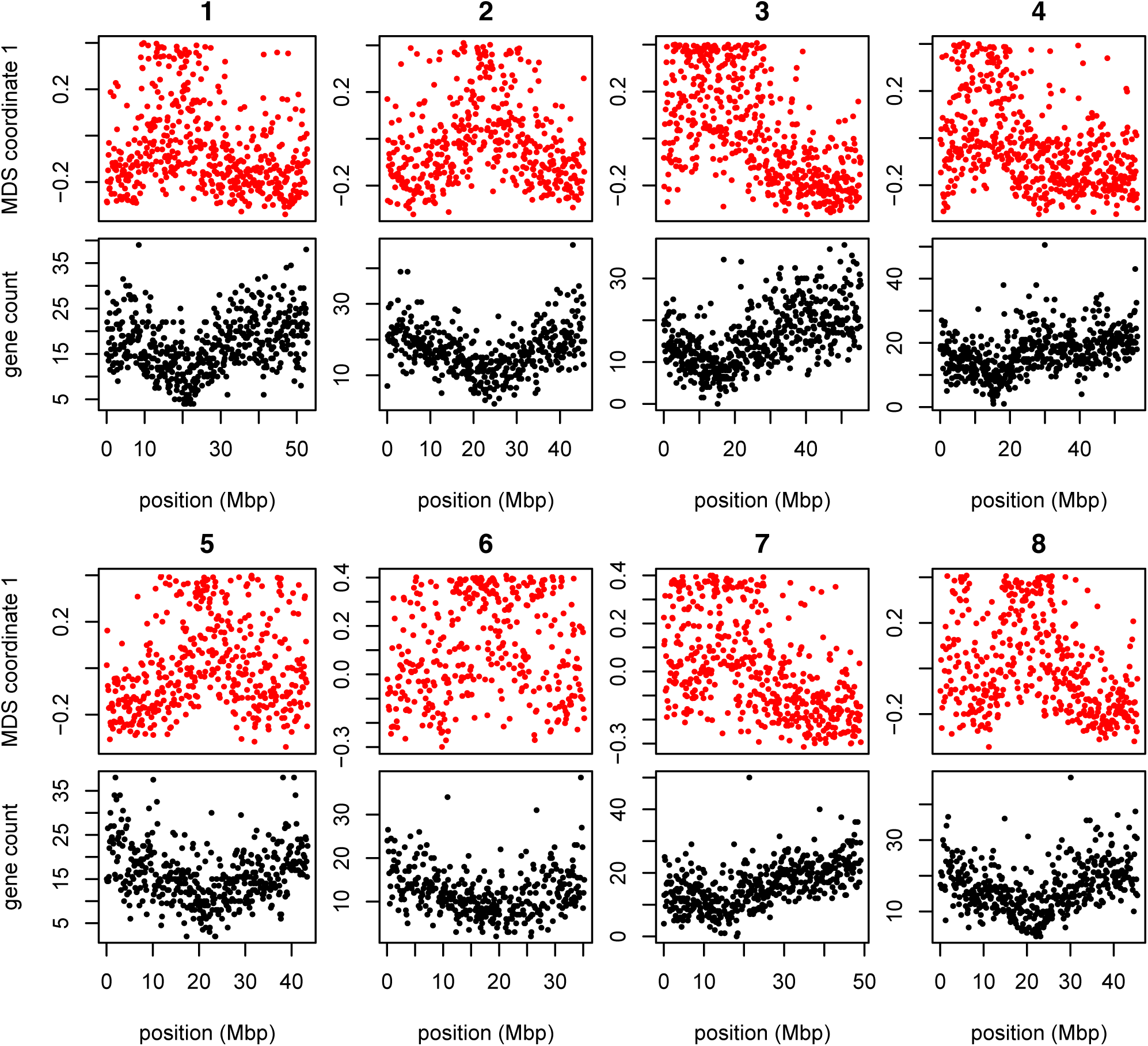
MDS coordinate and gene density for each window in the Medicago genome, for chromosomes 1–8 (numbered above each pair of figures). For each chromosome, the red plot above is first coordinate of MDS against the middle position of each window along each chromosome. The black plot below is gene count for each window against the middle position of each window.

The results were highly consistent across window sizes, window types (SNPs or bp), and number of PCs, as shown in Table S2.

## 4 Discussion

Our investigations have found substantial variation in the patterns of relatedness formed by population structure across the genomes of three diverse species, revealing distinct biological processes driving this variation in each species. More investigation, particularly on more species and datasets, will help to uncover what aspects of species history can explain these differences. With growing appreciation of the heterogeneous effects of selection across the genome, especially the importance of adaptive introgression and hybrid speciation [5, 23, 31, 57, 62], local adaptation [41, 66], and inversion polymorphisms [37, 38], local PCA may prove to be a useful exploratory tool to discover important genomic features.

We now discuss possible implications of this variation in the effects of population structure, the impact of various parameter choices in implementing the method, and possible additional applications.

**Chromosomal inversions** A major driver of variation in patterns of relatedness in two datasets we examined seems to be inversions. This may be common, but the example of *Medicago truncatula* shows that polymorphic inversions are not ubiquitous. PCA has been proposed as a method for discovering inversions [43]; however, the signal left by inversions likely cannot be distinguished from long haplotypes under balancing selection or simply regions of reduced recombination without additional lines of evidence. Inversions show up in our method because across the inverted region, most gene trees share a common split that dates back to the origin of the inversion. However, in many applications, inversions are a nuisance. For instance, SMARTPCA [54] reduces their effect on PCA plots by regressing out the effect of linked SNPs on each other. Removing samples with the less common orientation of each inversion reduced, but did not eliminate, the signal of inversions seen in the *Drosophila melanogaster* dataset, demonstrating that the genomic effects of transiently polymorphic inversions may outlast the inversions themselves.

**The effect of selection** Neutral processes are not expected to produce the chromosome-scale correlations we see in patterns of relatedness in the *Medicago truncatula* and *Drosophila melanogaster* datasets, because correlations induced by neutral processes should extend no further than does linkage disequilibrium (i.e., much less than a chromosome’s length). This suggests that they are produced by linked selection, a hypothesis backed up by correlations with gene density and recombination rate. We have also shown with simulations that linked selection can, in at least some circumstances, produce the sorts of patterns we observe. How might selection cause variation in patterns of relatedness? For instance, background selection (the effect on linked sites of selection against deleterious mutations Charlesworth et al. [10], Charlesworth [13]), can informally be thought of as reducing the number of potential contributors to the gene pool in regions of the genome with many possible deleterious mutations [29]. For this reason, if it acts in a spatial context, it is expected to induce samples from nearby locations to cluster together more frequently. Therefore, regions of the genome harboring many targets of local adaptation may show similar patterns, since migrant alleles in these regions will be selected against, and so locally gene trees will more closely reflect spatial proximity. Other forms of selection, such as hard sweeps on new mutations, repeated selection on standing variation, local adaptation, or temporally fluctuating selection, could clearly lead to variation in geographic patterns of relatedness in a similar way.

Another possible contributor is recent admixture between previously separated populations, the effects of which were not uniform across the genome due to selection. For instance, it has been hypothesized that large-scale variation in amount of introgressed Neanderthal DNA along the genome is due to selection against Neanderthal genes, leading to greater introgression in regions of lower gene density [27, 33]. African *Drosophila melanogaster* are thought to have a substantial amount of recently introgressed genome from “cosmopolitan” sources; if selection regularly favors genes from one origin, this could lead to substantial variation in patterns of relatedness correlated with local gene density.

There has been substantial debate over the relative impacts of different forms of selection (e.g., Burri et al. [8], Charlesworth et al. [11], Charlesworth [12], Corbett-Detig et al. [17], Harris and Nielsen [27], Hedrick [28], Martin et al. [46], Pease and Hahn [55], Phung et al. [56], Stankowski et al. [61]). These have been difficult to disentangle in part because for the most part theory makes predictions which are only strictly valid in randomly mating (i.e., unstructured) populations, and it is unclear to what extent the spatial structure observed in most real populations will affect these predictions. It may be possible to design more powerful statistics that make stronger use of spatial information.

**Parameter choices** There are several choices in the method that may in principle affect the results. As with whole-genome PCA, the choice of samples is important, as variation not strongly represented in the sample will not be discovered. The effects of strongly imbalanced sampling schemes are often corrected by dropping samples in overrepresented groups; but downweighting may be a better option that does not discard data. Next, the choice of window size may be important, although in our applications results were not sensitive to this. Finally, which collections of genomic regions are compared to each other (steps 3 and 4 in Figure 1), along with the method used to discover common structure, will affect results. We used MDS, applied to either each chromosome separately or to the entire genome; for instance, human inversions are clearly visible as outliers when compared to the rest of their chromosomes, but genome-wide, their signal is obscured by the numerous other signals of comparable strength.

Besides window length, there is also the question of how to choose windows. In these applications we have used nonoverlapping windows with equal numbers of polymorphic sites. However, we found little change in results when using different window sizes or when measuring windows in physical distance (in bp).

Finally, our software allows different choices for how many PCs to use in approximating structure of each window (*k* in equation 1), and how many MDS coordinates to use when describing the distance matrix between windows, but in our exploration, changing these has not produced dramatically different results. These are all part of more general techniques in dimension reduction and high-dimensional data visualization; we encourage the user to experiment.

**Applications** So-called cryptic relatedness between samples has been one of the major sources of confounding in genome-wide association studies (GWAS) and so methods must account for it by modeling population structure or kinship [2, 69]. Modern “mixed model” methods [e.g. 42] account for this with either a single, genome-wide kinship matrix or one constructed using only sites unlinked to the focal SNP. Since the effects of population structure is not constant along the genome, this could in principle lead to an inflation of false positives in parts of the genome with stronger population structure than the genome-wide average. A method such as ours might be used to estimate local kinship matrices, thus providing a more sensitive correction, although doing so without removing the signal itself could be challenging. Fortunately, in our human dataset this does not seem likely to have a strong effect: most variation is due to small, independent regions, possibly primarily inversions, and so may not have a major effect on GWAS. In the other species we examined, particularly *Drosophila melanogaster*, treating population structure as a single quantity would entail a substantial loss of power, and could potentially be misleading.

## Acknowledgements

We are indebted to John Pool, Russ Corbett-Detig, Matilde Cordeiro, and Peter Chang for assistance with obtaining data and interpreting results (especially inversion status of *D*. *melanogaster* samples). Jaime Ashander and Jerome Kelleher provided assistance in performing the simulations. Thanks also go to Yaniv Brandvain, Barbara Engelhardt, Charles Langley, Graham Coop, and Jeremy Berg for helpful comments and for encouraging the project.

## A Choosing window length

The choice of window length entails a balance between signal and noise. In very short windows, genealogies of the samples will only be represented by a few trees, so variation between windows represents demographic noise rather than meaningful variation in patterns of relatedness. Longer windows generally have more distinct trees (and SNPs), allowing for less noisy estimation of local patterns of relatedness. However, to better resolve meaningful signal, i.e., differences in patterns of relatedness along the genome, we would like reasonably short windows.

Since we summarize patterns of relatedness using relative positions in the principal component maps, we quantify “noise” as the standard error of a sample’s position on PC1 in a particular window, averaged across windows and samples, and “signal” as the standard deviation of the sample’s position on PC1 over all windows, averaged over samples. The definition of eigenvectors does not specify their sign, and so when comparing between windows we choose signs to best match each other: after choosing *PC*1_1_, for instance, if *u* is the first eigenvector obtained from the covariance matrix for window *j*, then we next choose *PC*1_*j*_ = *±u*, where the sign is chosen according to which of ‖*PC*1_1_ - *u*‖ or ‖*PC*1_1_ + *u*‖ is smaller.

After doing this, the mean variance across windows is

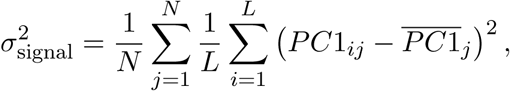

where *PC*1_*ij*_ is the position of the *i*^th^ individual on *PC*1 in window *j*, and 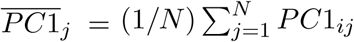. We estimate the standard error for each *PC*1_*ij*_ using the block jack-knife [9, 20]: we divide the *j*^th^ window into 10 equal-sized pieces, and let *PC*1_*ij,k*_ denote the first principal component of this region found after removing the *k*^th^ piece; then the estimate of the squared standard error is 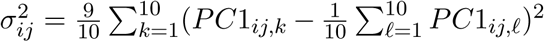. Averaging over samples and windows,

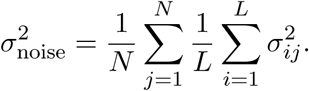

For the main analysis, we defined windows to each consist of the same number of neighboring SNPs, and calculated 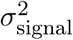 and 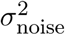 for a range of window sizes (i.e., numbers of SNPs). For our main results we chose the smallest window for which 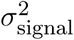 was consistently larger than 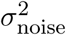 (but checked other sizes); the values for various window sizes across *Drosophila* chromosomes are shown in Table S1. In the cases we examined, we found nearly identical results after varying window size, and choosing windows to be of the same physical length (in bp) rather than in numbers of SNPs.

**Table S1:** Measures of signal and noise, computed separately for each chromosome arm in the *Drosophila* dataset, at different window sizes. All values are multiplied by 1, 000. Starting at windows of 1,000 SNPs, the signal (variation of PC1 between windows) starts to be substantially larger than the noise (standard error of PC1 for each window).

| chrom. arm |  | window length (SNPs) |  |  |  |  |
| --- | --- | --- | --- | --- | --- | --- |
|  |  | 100 | 500 | 1,000 | 10,000 | 100,000 |
| 2L | $\sigma_{\text{noise}}^2$ | 2.05 | 1.64 | 1.18 | 0.17 | 0.04 |
| | $\sigma_{\text{signal}}^2$ | 2.76 | 2.69 | 2.23 | 0.68 | 0.31 |
| 2R | $\sigma_{\text{noise}}^2$ | 2.18 | 1.92 | 1.63 | 0.58 | 0.13 |
| | $\sigma_{\text{signal}}^2$ | 2.78 | 2.70 | 2.65 | 2.31 | 1.82 |
| 3L | $\sigma_{\text{noise}}^2$ | 2.08 | 2.00 | 1.64 | 0.73 | 0.25 |
| | $\sigma_{\text{signal}}^2$ | 2.60 | 2.52 | 2.40 | 1.68 | 1.89 |
| 3R | $\sigma_{\text{noise}}^2$ | 1.95 | 1.76 | 1.44 | 0.59 | 0.20 |
| | $\sigma_{\text{signal}}^2$ | 2.58 | 2.51 | 2.44 | 1.96 | 1.40 |
| X | $\sigma_{\text{noise}}^2$ | 2.48 | 2.04 | 1.54 | 1.62 | 0.17 |
| | $\sigma_{\text{signal}}^2$ | 2.61 | 2.43 | 2.30 | 0.32 | 1.14 |

## B Simulations

We implemented two types of simulation: first, simple simulations of Gaussian “genotypes” where the expectation of variation in “population structure” was clear; and next, individual-based simulations with explicit genomes, using SLiM.

### B.1 Gaussian simulations

We simulated genotypes at each locus independently, drawing each vector of genotypes from a multivariate Gaussian distribution with zero mean and covariance matrix Σ. Sampled individuals came from three populations, and each Σ_*ij*_ depends on which populations the individuals *i* and *j* are in, as well as the location along the chromosome. There are three population-level mean relatedness matrices along the genome, which apply to the first quarter (*S*^(1)^), the middle half (*S*^(2)^), and the last quarter (*S*^(3)^), respectively:

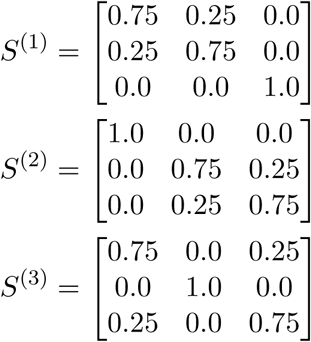

If indiviudals *i* and *j* are in populations *p*(*i*) and *p*(*j*) respectively, then the covariance between their genotypes is Σ_*ij*_ = *S*_*p*(*i*),*p*(*j*)_, using the appropriate *S* for that segment of the genome. The variance of individual *i*’s genotype is Σ_*ii*_ = *S*_*p*(*i*),*p*(*i*)_ + 0.1.

We first created “genotypes” in this way with fifty individuals from each of the three populations; running our method on a genome with 99 windows of 400 loci each produced the first plot in Figure S4. These matrices are chosen so that the top two eigenvalues Σ are the same (both 50.1), and so the ordering of the top two PCs is arbitrary. If our method was sensitive to PC ordering, then half the windows in each region that have one ordering would cluster with each other, separate from the other half.

We then marked each genotype in the first half of the chromosome as missing, independently, with probability 1/2 and ran our method again, producing the second plot of Figure S4. If our method was influenced by missing data, we would expect the first half of the chromosome to separate from the second in the MDS plot.

### B.2 SLiM simulations

Our SLiM simulations were constructed as follows. Individuals are diploid, and genomes have a length of 153,520,244 bp. Recombination was either (a) flat, with a constant rate of 10^−9^; (b) according to the human female HapMap map for chromosome 7; or (c) constant in each of seven equal-sized regions, beginning at 2.04×10^−8^, descending by a factor of four for three steps, and then ascending by a factor of four for three steps, so that the middle seventh has the lowest recombination rate, and the outer two sevenths has a rate 64 times higher. Selected mutations are introduced at a rate of 10^−10^ per bp per individual per generation, and have selection coefficients drawn from a Gamma distribution with mean 0.005 and shape 2; each coefficient are either positive or negative with probabilities 1/30 and 29/30 respectively. Each simulation was run for 50,000 generations.

Each individual has a spatial position in the two-dimensional square of width *W* = 8. Each time step, each individual chooses the nearest other to mate with, producing a random, Poisson distributed number of offspring with mean 1/3. Offspring are assigned random spatial locations displaced from their parent’s by a bivariate Gaussian with mean zero and standard deviation *σ* = 0.2, reflected to stay within the habitat range.

Each individual survives to the next time step with probability equal to their fitness. Fitness values are determined multiplicatively by the effects of each mutation, but are multiplied by an additional factor determined by the local density of individuals. This factor is equal to *ρ*/(1 + *C*), where *ρ* = 2*πKσ*^2^ is the carrying capacity per circle of radius *σ*; *K* = 100 is the mean equilibrium population density; and *C* is the sum of a Gaussian kernel with standard deviation *σ* = 0.1 between the focal individual and all other individuals within distance 3*σ*. To avoid edge effects, fitnesses are further multiplied by min(1, *z*), where *z* is the distance to the nearest boundary. This produces populations that fluctuate at equilibrium around 6,000 individuals in total, fairly evenly spread across the square.

In one additional simulation, we modified fitnesses by multiplying the selective effect of each allele in each individual by multiplying it by 2*x*/*W* - 1, where *x* is the *x* coordinate of the individual. This makes the effect of each allele opposite on the left and on the right, and neutral in the middle, and leads to a moderate number of balanced polymorphisms.

## C Supplementary Tables

## D Supplementary Figures

**Figure S1:**
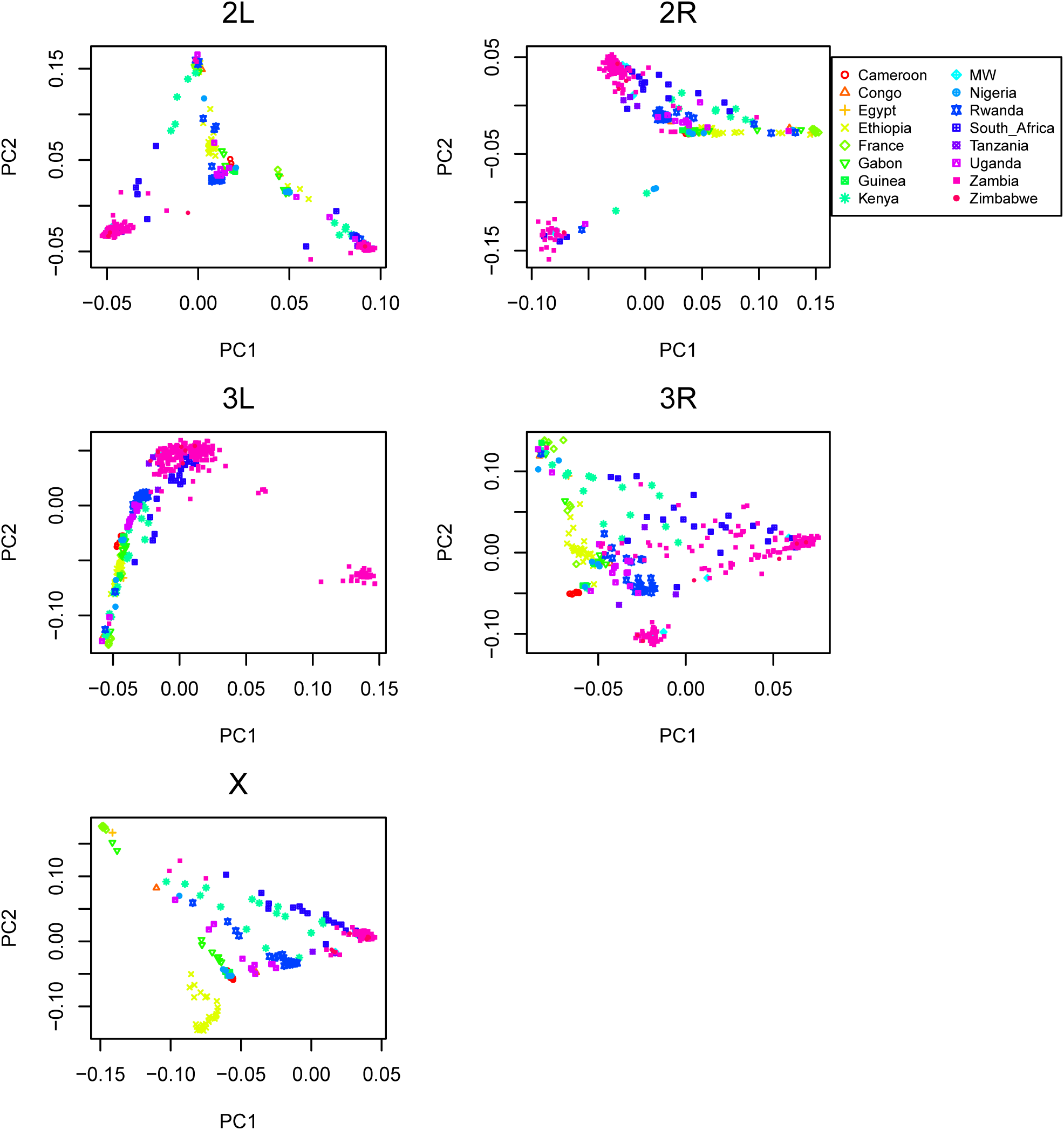
PCA plots for chromosome arms 2L, 2R, 3L, 3R and X of the *Drosophila melanogaster* dataset.

**Figure S2:**
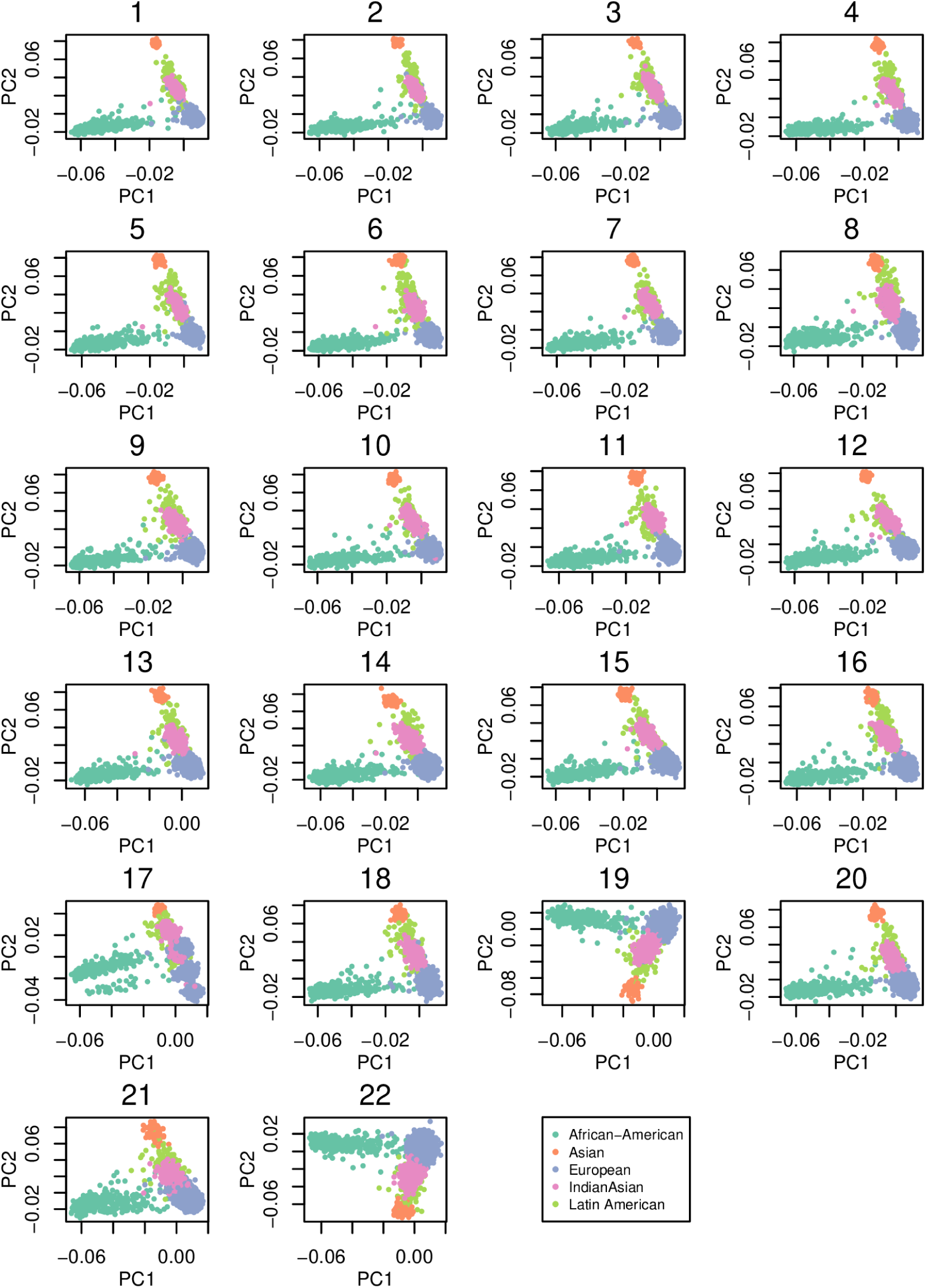
PCA plots for all 22 human autosomes from the POPRES data.

**Table S2:**
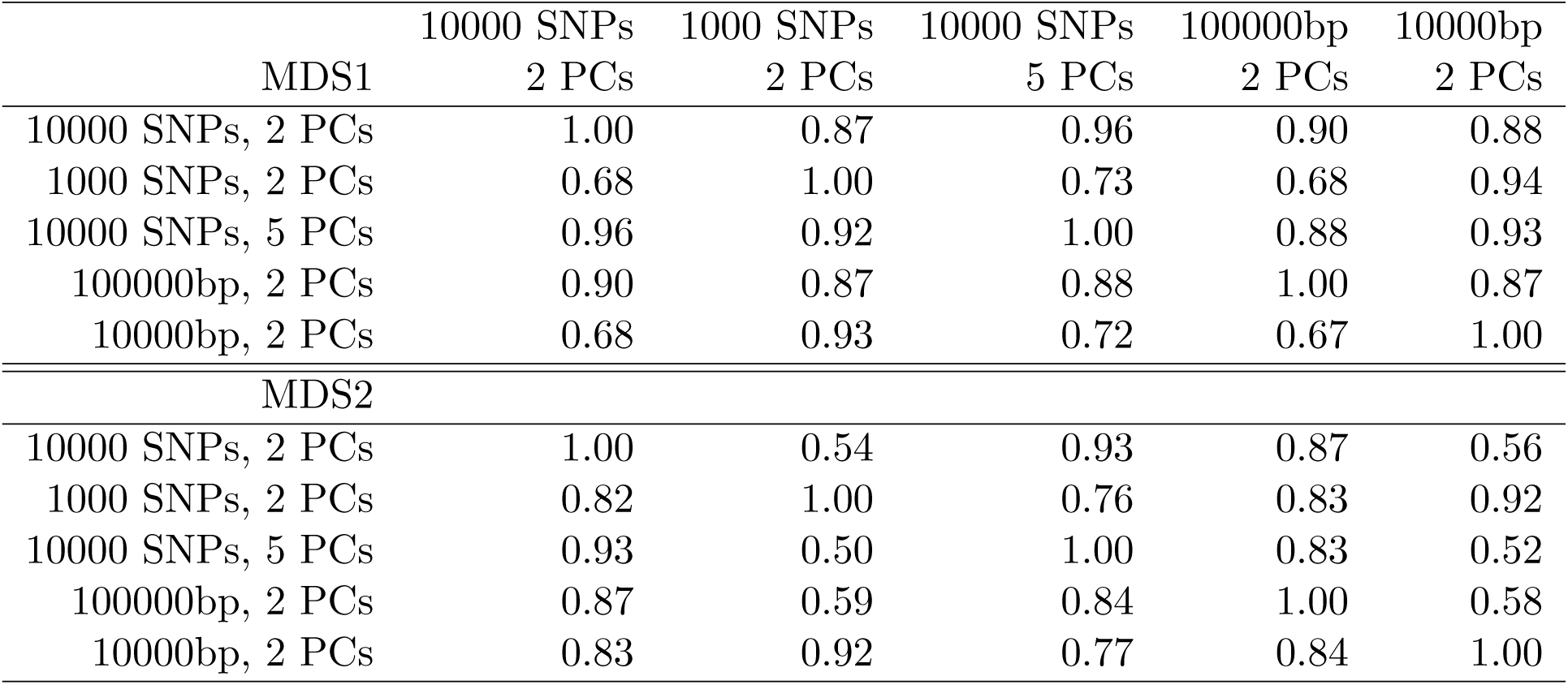
Correlations between MDS coordinates of genomic regions between runs with different parameter values. To produce these, we first ran the algorithm with the specified window size and number of PCs (*k* in equation (1)) on the full *Medicago truncatula* dataset. Then to obtain the correlation between results obtained from parameters A in the row of the matrix above and parameters B in the column of the matrix above, we mapped the windows of B to those of A by averaging MDS coordinates of any windows of B whose midpoints lay in the corresponding window of A; we then computed the correlation between the MDS coordinates of A and the averaged MDS coordinates of B. This is not a symmetric operation, so these matrices are not symmetric. As expected, parameter values with smaller windows produce noisier estimates, but plots of MDS values along the genome are visually very similar.

**Figure S3:**
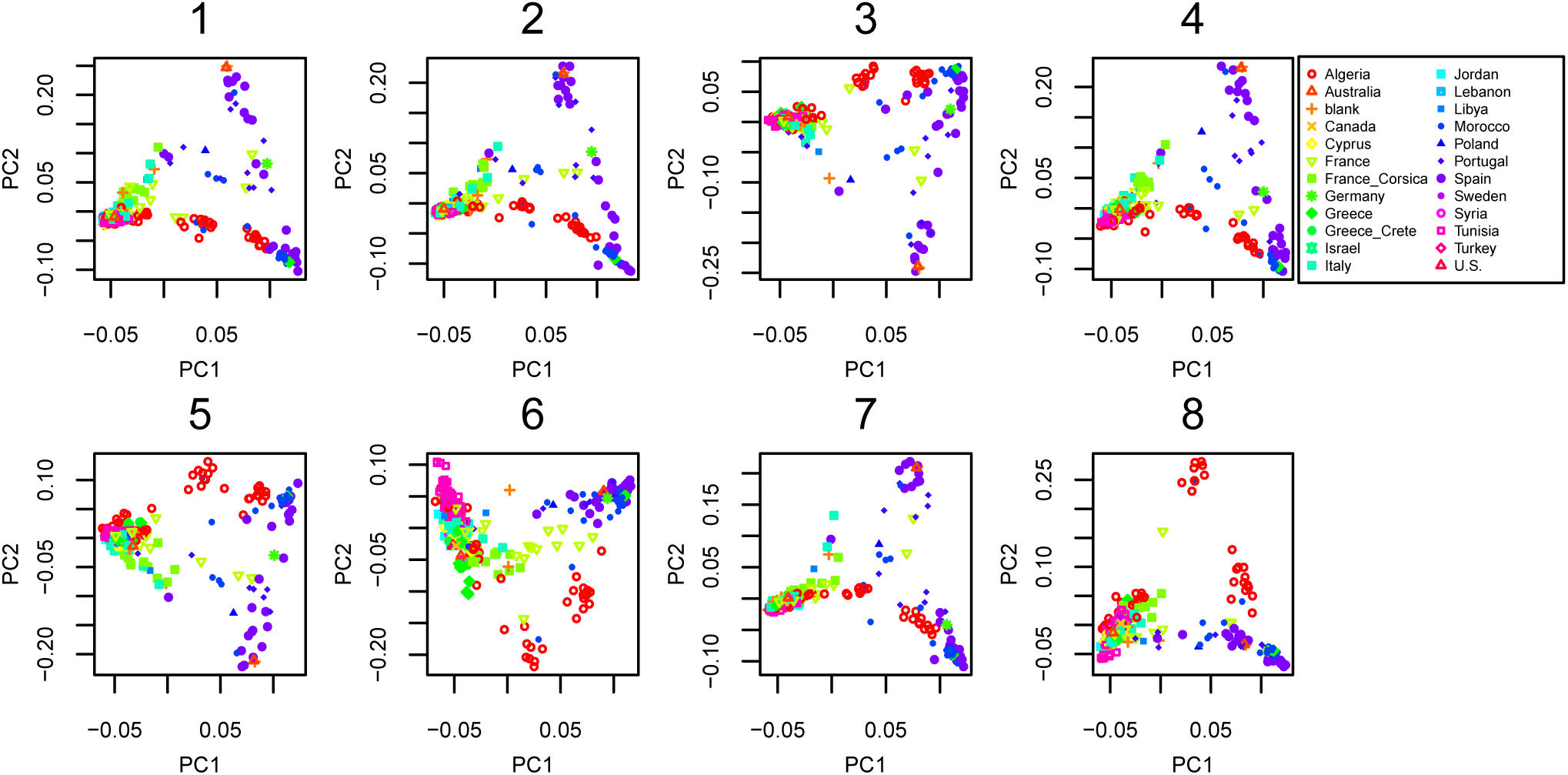
PCA plots for all 8 chromosomes in the *Medicago truncatula* dataset.

**Figure S4:**
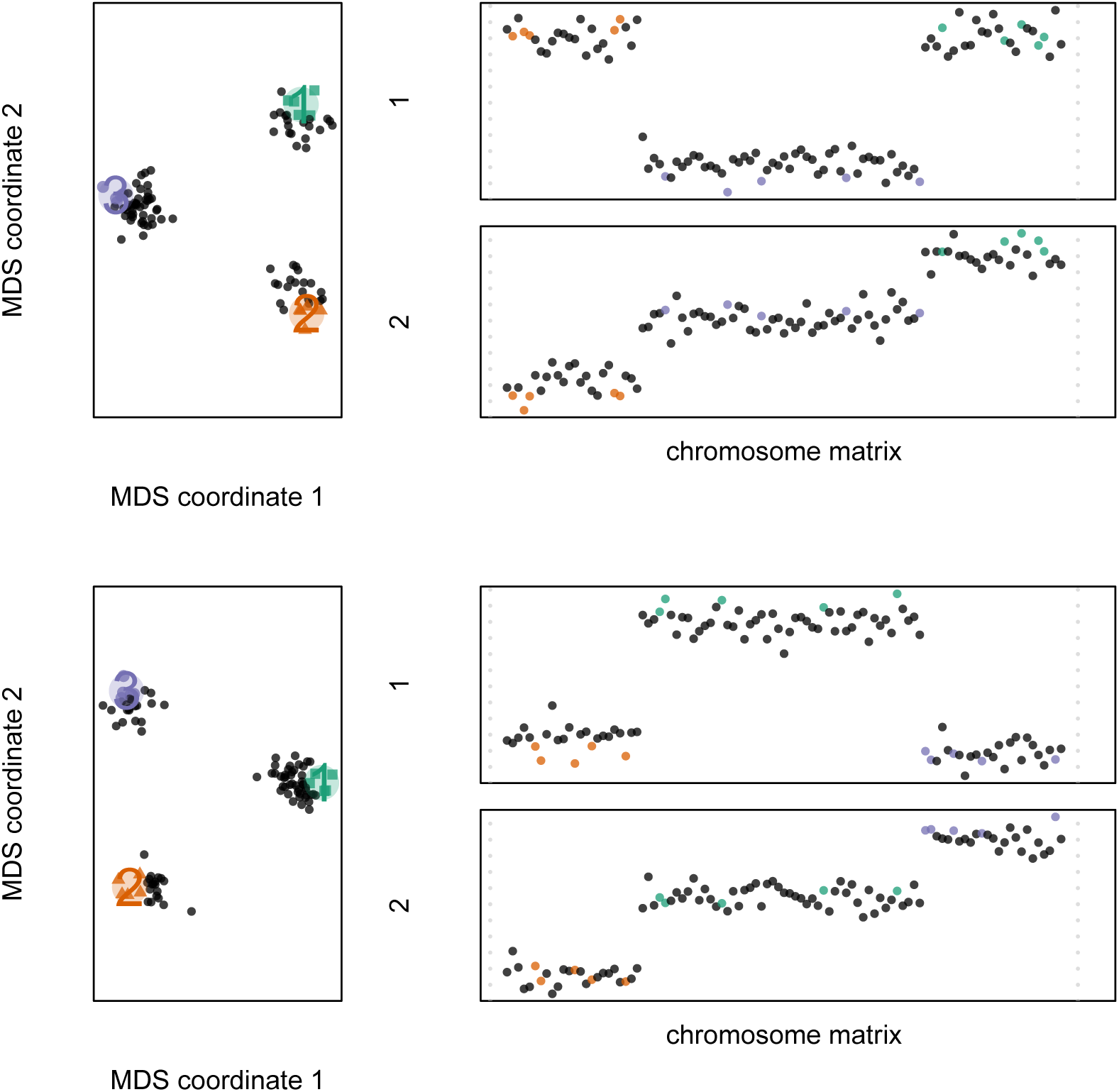
MDS visualizations of the Gaussian genotypes described in Appendix B.1, for 50 individuals from each of three populations. **(top)** The first quarter, middle half, and final quarter of the chromosome each have different population structure, as expected, despite the possibility for PC switching within each. **(bottom)** The same picture results even after marking a random 50% of the genotypes in the first half of the chromosome as missing.

**Figure S5:**
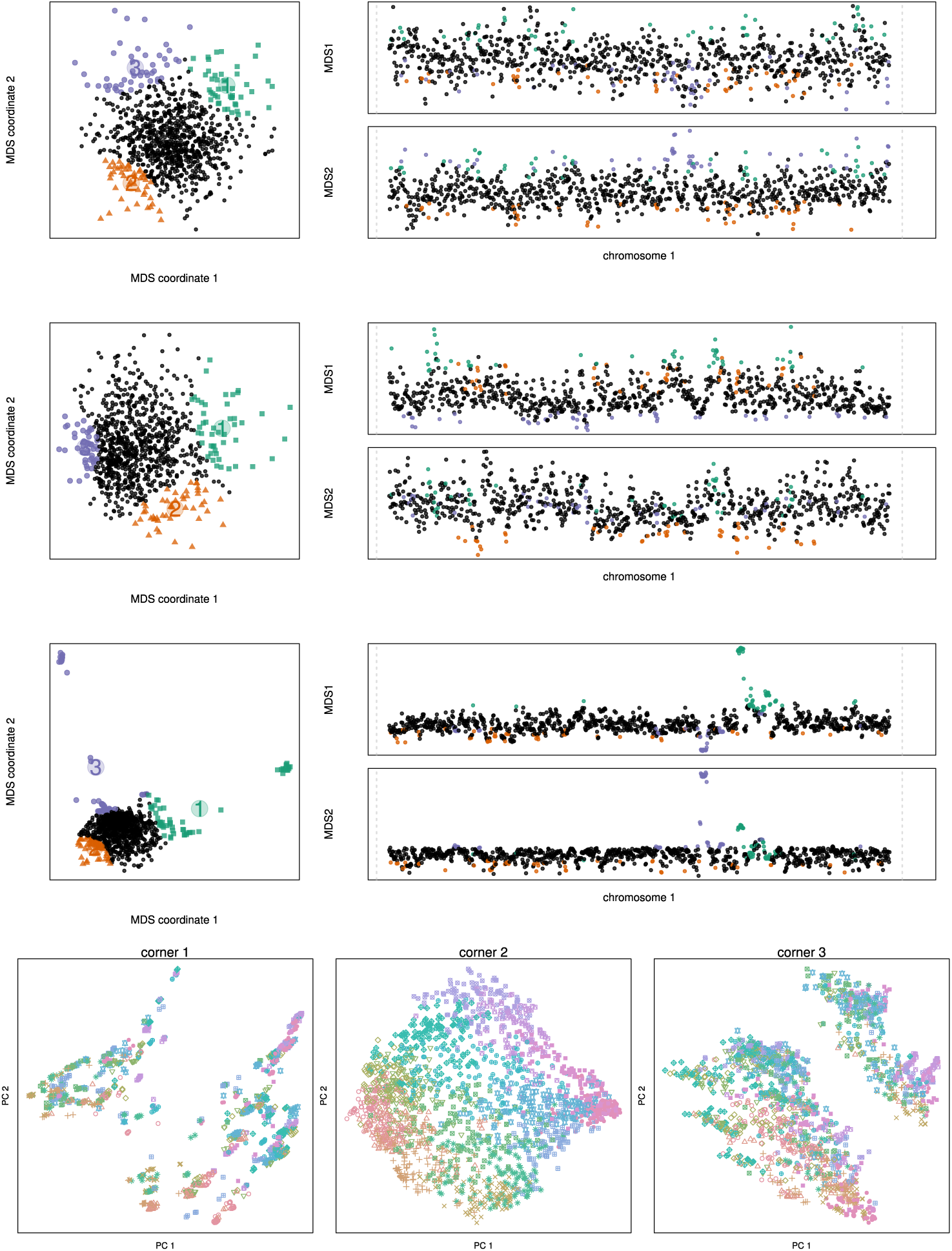
MDS visualizations of the results of individual-based simulations using SLiM (see Appendix B.2 for details). All simulations are neutral, and recombination is: **(top)** constant; **(top middle)** varies stepwise by factors of two in seven equal-length segments, with highest rates on the ends, so the middle segment has a recombination rate 64 times lower than the ends; **(bottom middle)** according to the HapMap human female chromosome 7 map. The **bottom** figure shows PCA maps corresponding to the three colored windows of the last (HapMap) situation; the outlying regions are long regions of low recombination rate, so that region can be dominated by a few correlated trees, similar to an inversion.

**Figure S6:**
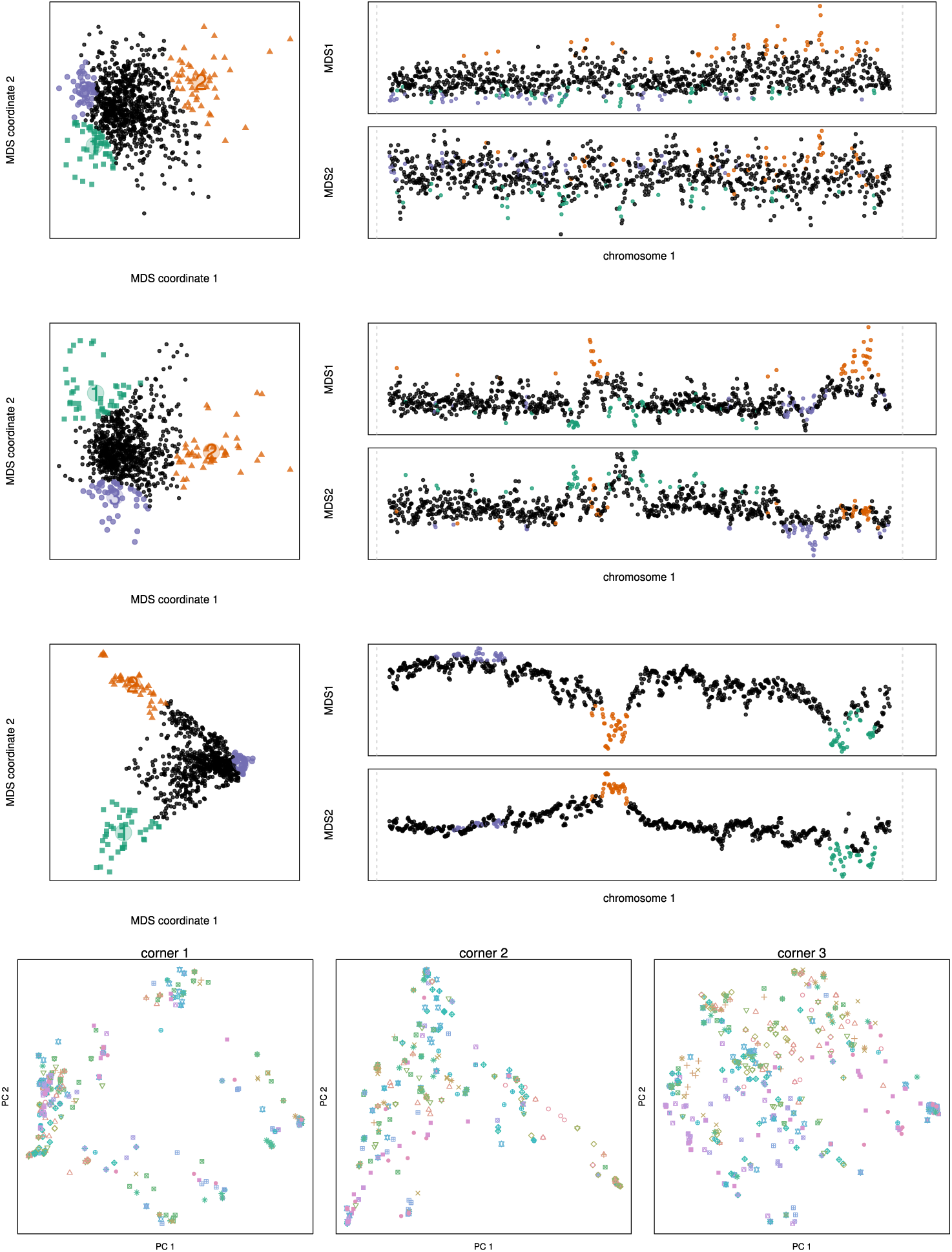
MDS visualizations of the results of individual-based simulations using SLiM (see Appendix B.2 for details). All simulations incorporate linked selection by allowing selected mutations to appear in the same two regions of the genome: the one-sixth of the genome immediately before the halfway point, and the last one-sixth of the genome. **(top)** Constant recombination rate. **(top middle)** Stepwise varying recombination rate (as described in Figure S5). **(bottom middle)** Constant recombination rate with spatially varying effects of selection. **(bottom)** PCAp lots corresponding to the highlighted corners of the last MDS visualization, showing how spatially varying linked selection has affected patterns of relatedness.

**Figure S7:**
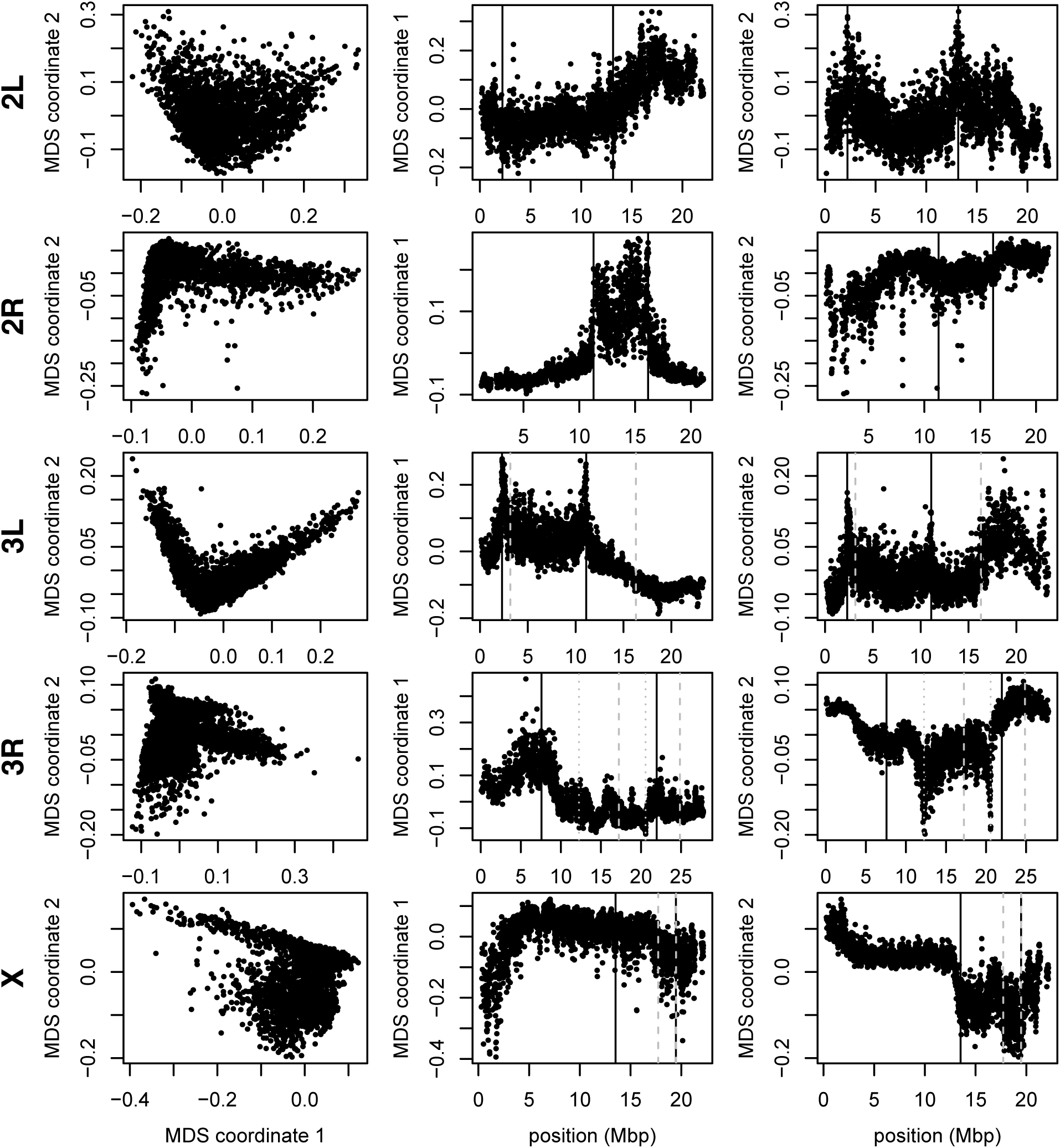
MDS visualizations for each chromosome arm of *Drosophila melanogaster*, as in Figure 2, except that the method was run using five PCs (*k* = 5) instead of two.

**Figure S8:**
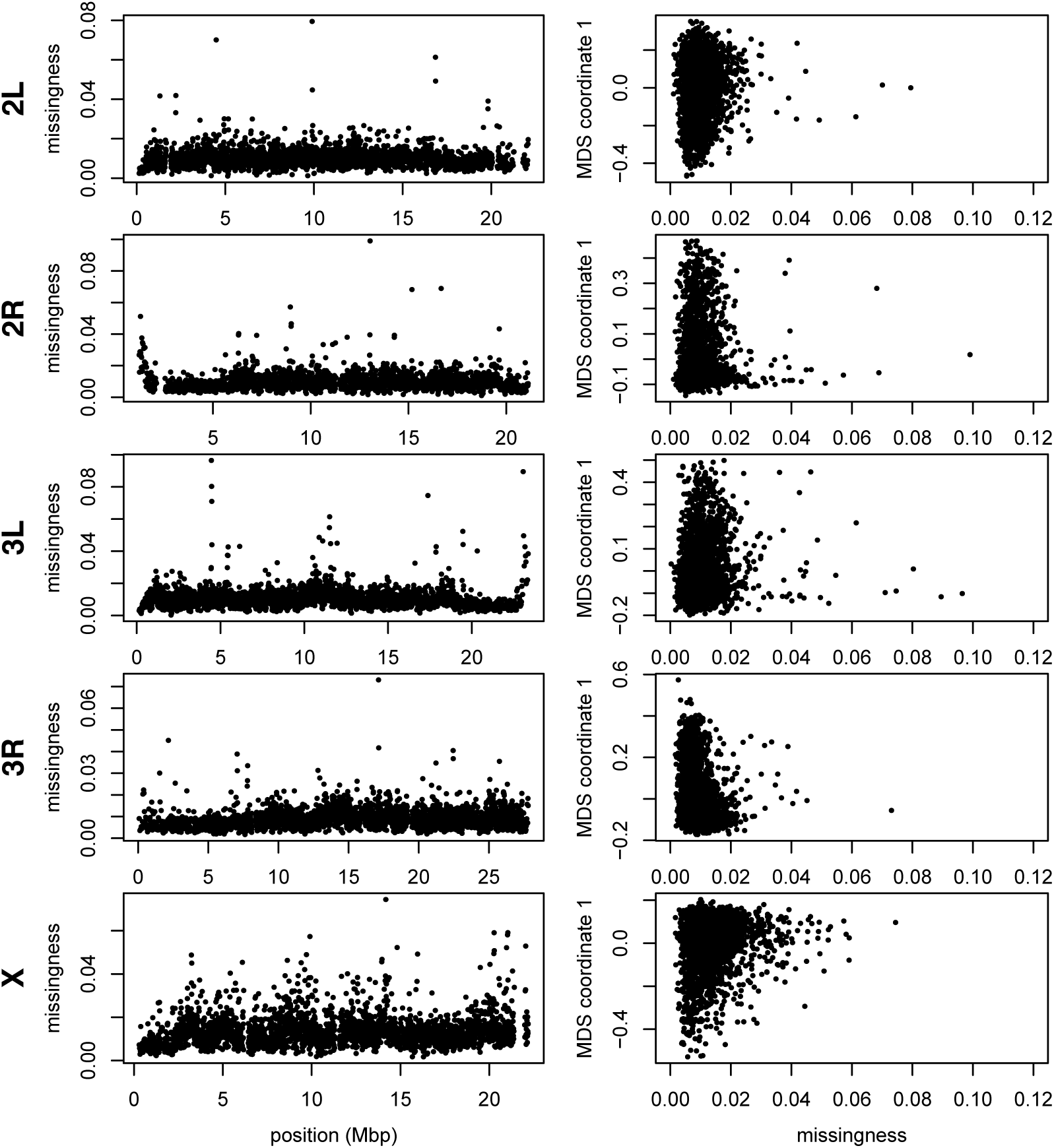
The proportion of data in each window that are missing, compared to the value of the first MDS coordinate for the *Drosophila melanogaster* data from Figure 2.

**Figure S9:**
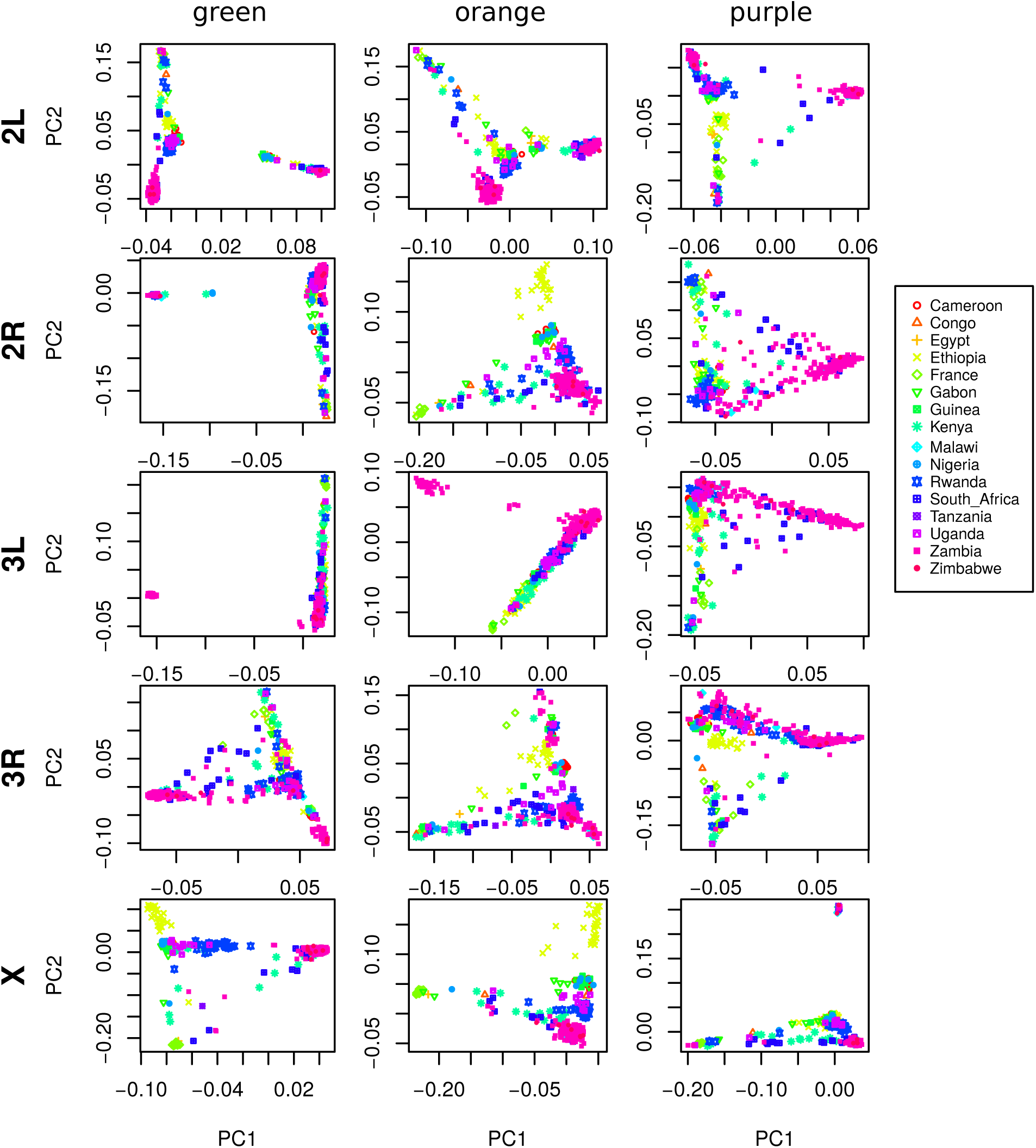
PCA plots for the three sets of genomic windows colored in Figure 2, on each chromosome arm of *Drosophila melanogaster*. In all plots, each point represents a sample. The first column shows the combined PCA plot for windows whose points are colored green in Figure 2; the second is for orange windows; and the third is for purple windows.

**Figure S10:**
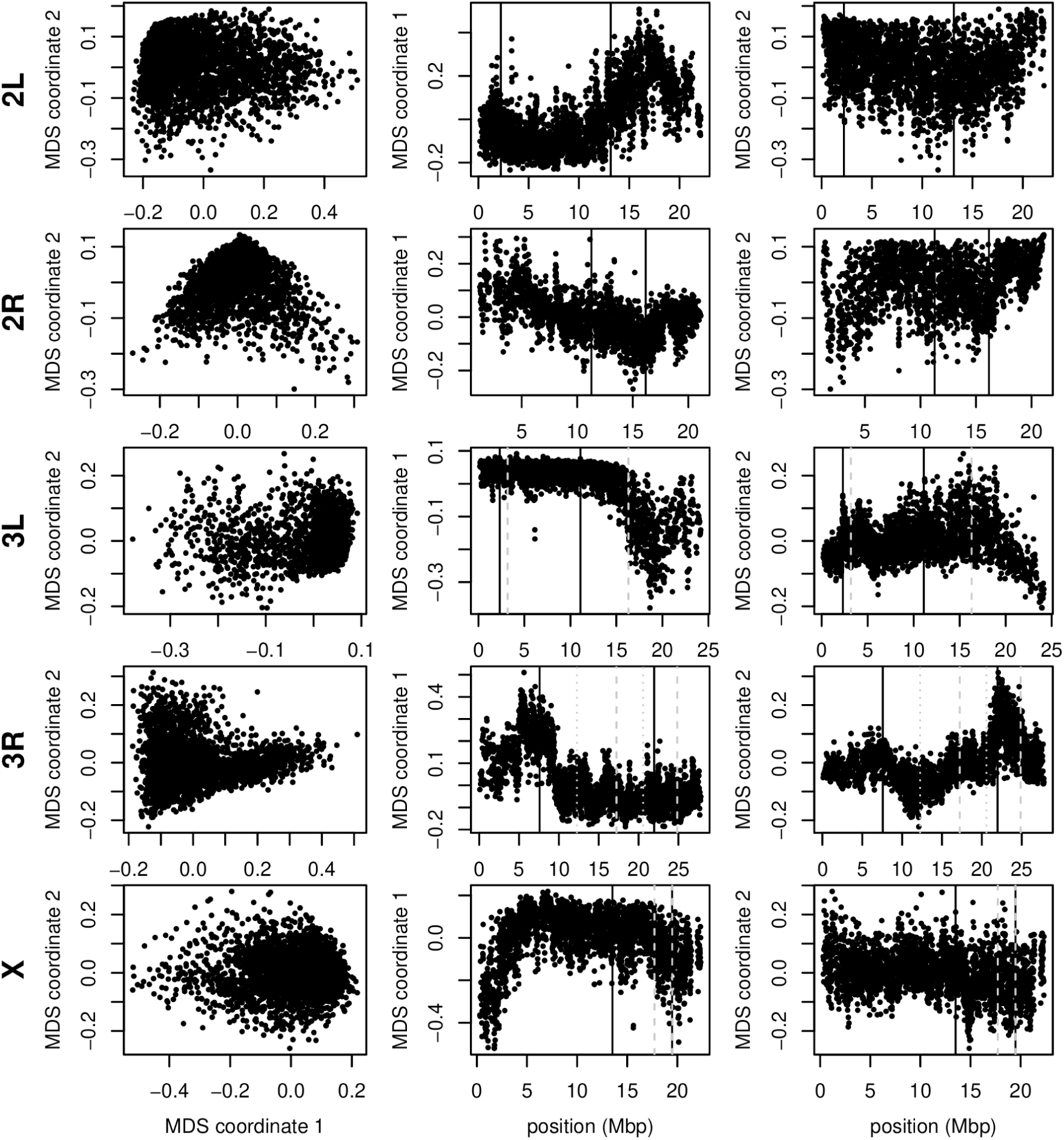
Variation in structure for windows of 1,000 SNPs across *Drosophila melanogaster* chromosome arms: without inversions. As in Figure 2, but after omitting for each chromosome arm individuals carrying the less frequent orientation of any inversions on that chromosome arm. The values differ from those in 4 in the window size used and that some MDS values were inverted (but relative orientation is meaningless as chromosome arms were run separately, unlike for *Medicago*). In all plots, each point represents one window along the genome. The first column shows the MDS visualization of relationships between windows, and the second and third columns show the midpoint of each window against the two MDS coordinates; rows correspond to chromosome arms. Vertical lines show the breakpoints of known polymorphic inversions.

**Figure S11:**
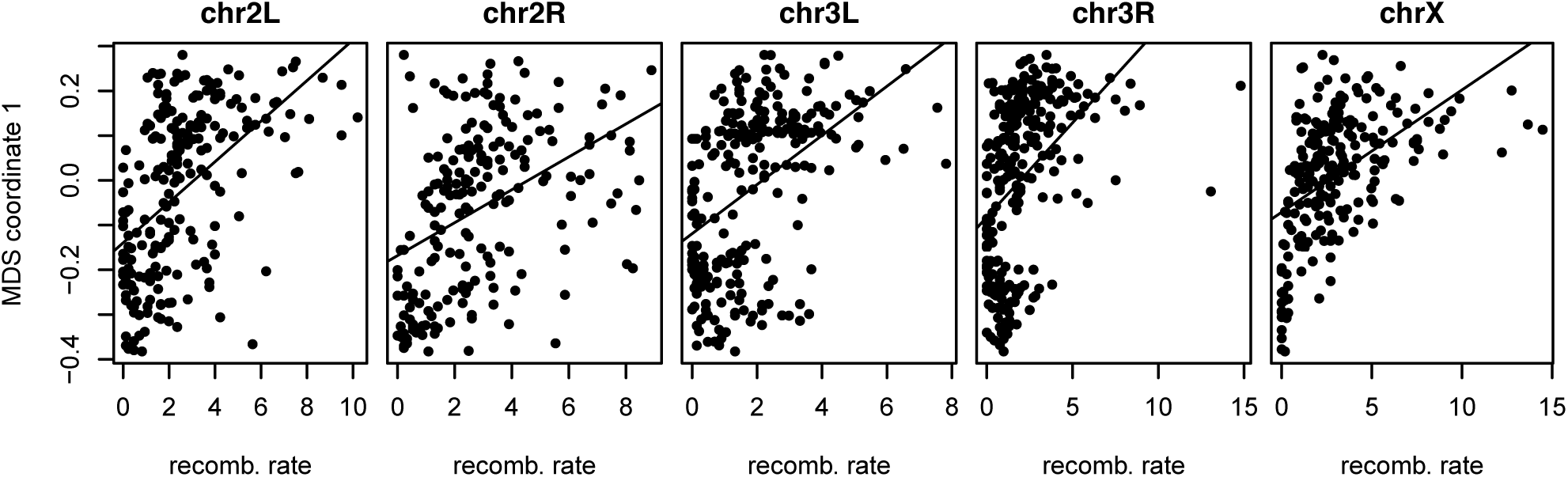
Recombination rate, and the effects of population structure for *Drosophila melanogaster*: this shows the first MDS coordinate and recombination rate (in cM/Mbp), as in Figure 4, against each other. Since the windows underlying estimates of Figure 4 do not coincide, to obtain correlations we divided the genome into 100Kbp bins, and for each variable (recombination rate and MDS coordinate 1) averaged the values of each overlapping bin with weight proportional to the proportion of overlap. The correlation coefficient and *p*-values for each linear regression are as follows: 2L: correlation = 0.52, *r*^2^ = 0.27; 2R: correlation = 0.43, *r*^2^ = 0.18; 3L: correlation = 0.47, *r*^2^ = 0.21; 3R: correlation = 0.46, *r*^2^ = 0.21; X: correlation = 0.50, *r*^2^ = 0.24.

**Figure S12:**
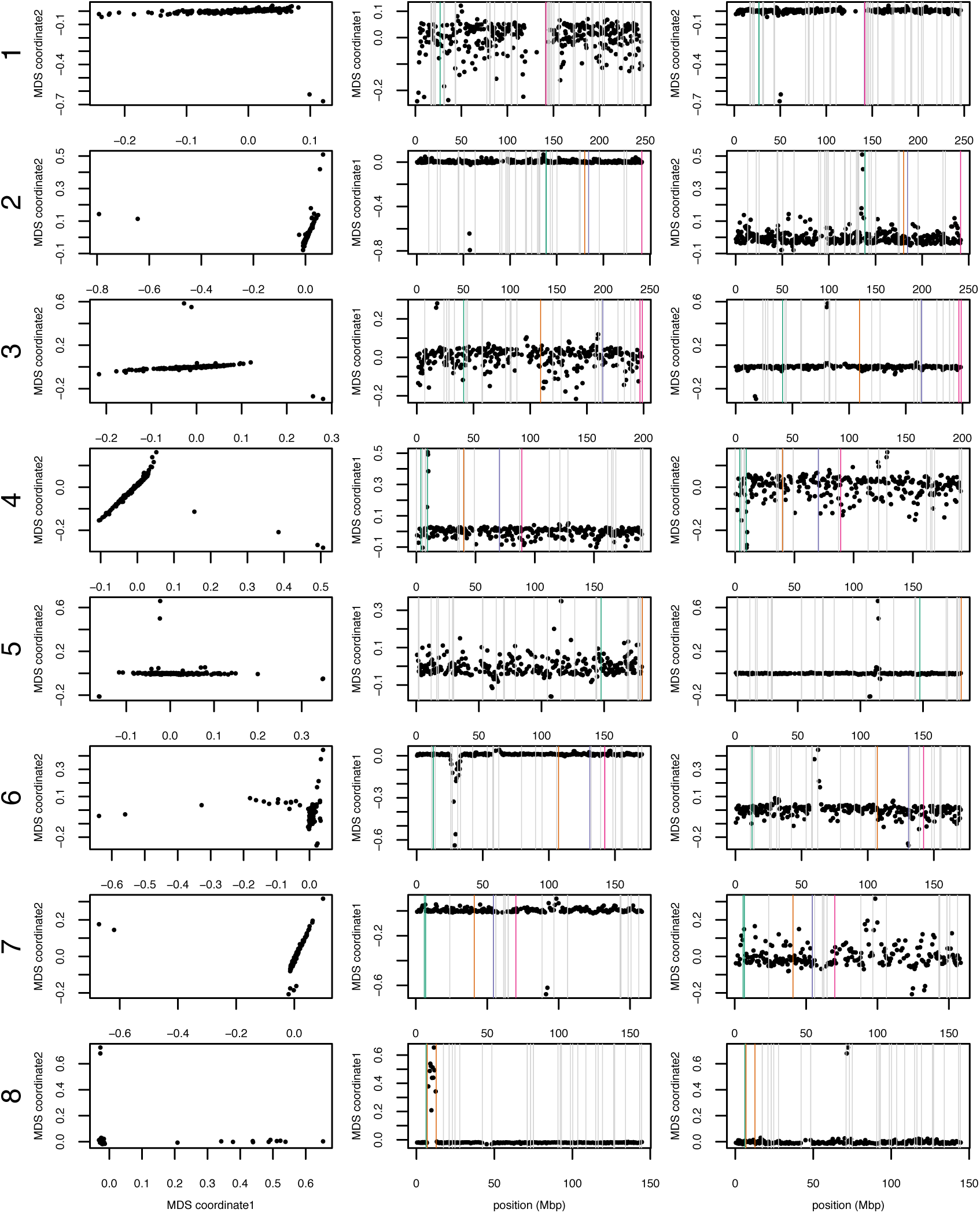
MDS plots for human chromosomes 1-8. The first column shows the MDS visualization of relationships between windows, and the second and third columns show the midpoint of each window against the two MDS coordinates; rows correspond to chromosomes. Colorful vertical lines show the breakpoints of known valid inversions, while grey vertical lines show the breakpoints of predicted inversions.

**Figure S13:**
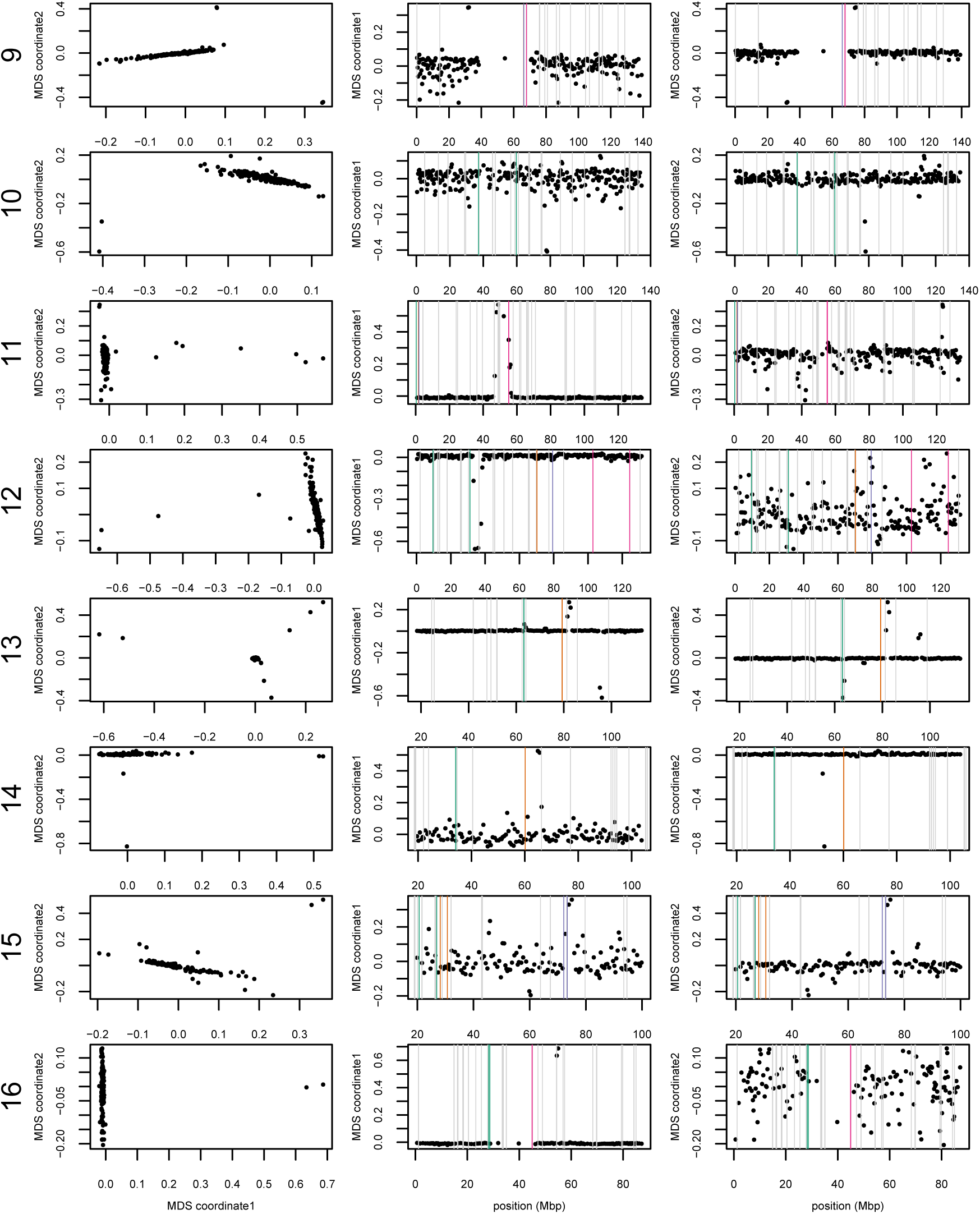
MDS plots for human chromosomes 9-16, as in Supplemental Figure S12.

**Figure S14:**
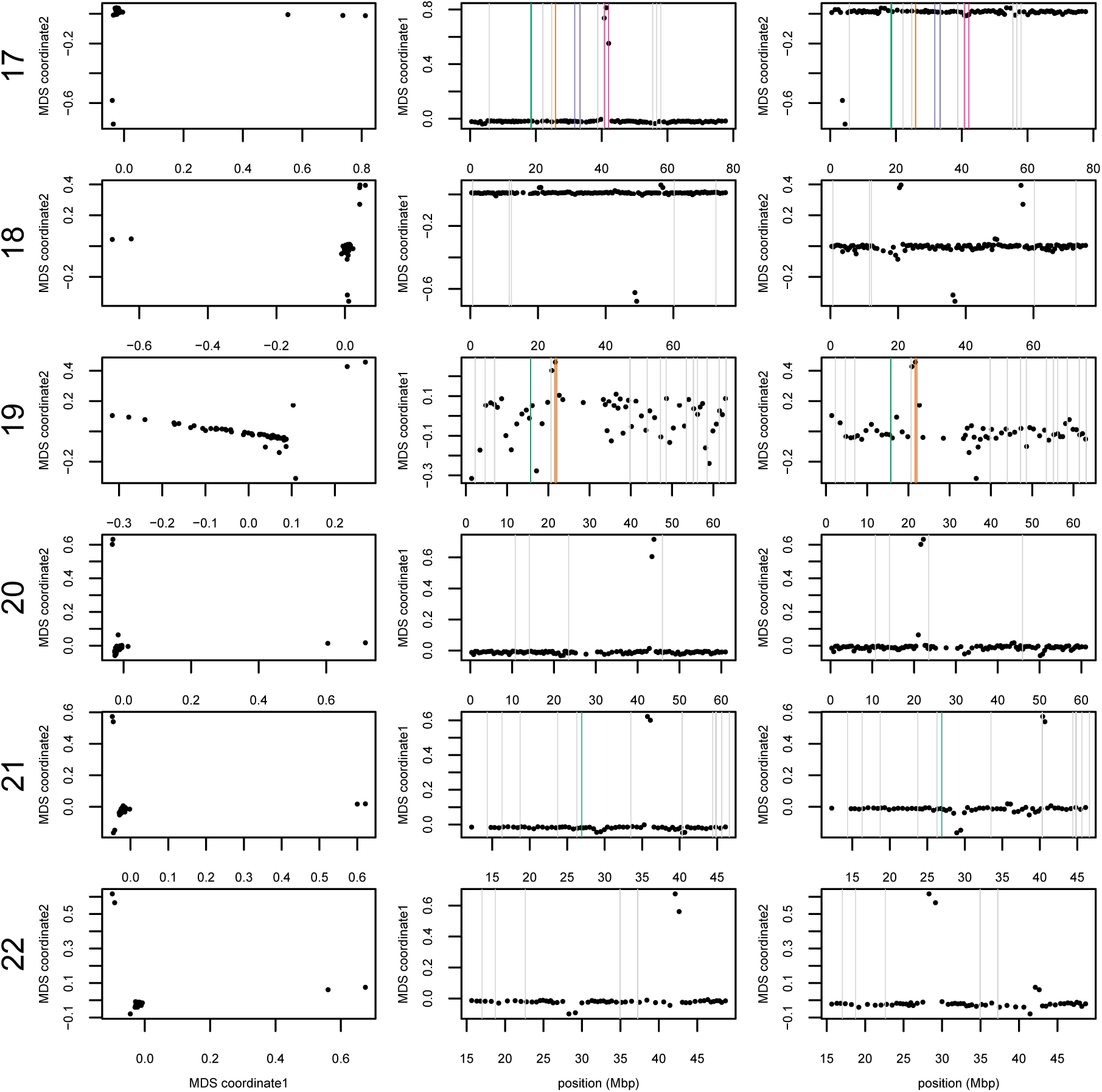
MDS plots for human chromosomes 17-22, as in Supplemental Figure S12.

**Figure S15:**
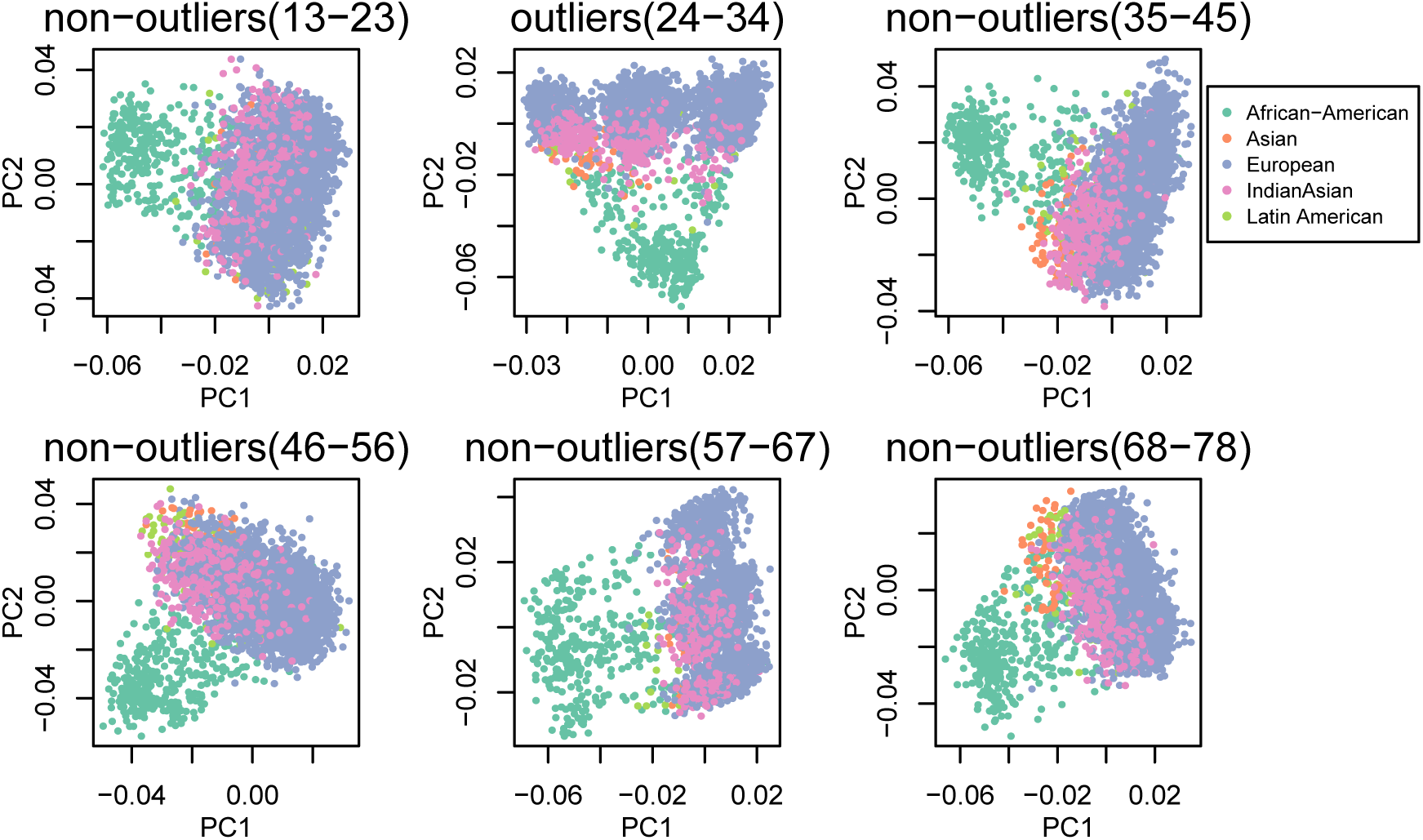
Comparison of PCA figures within outlying windows (center column) and flanking non-outlying windows (left and right columns) for the two windows having outlying MDS scores on chromosome 8.

**Figure S16:**
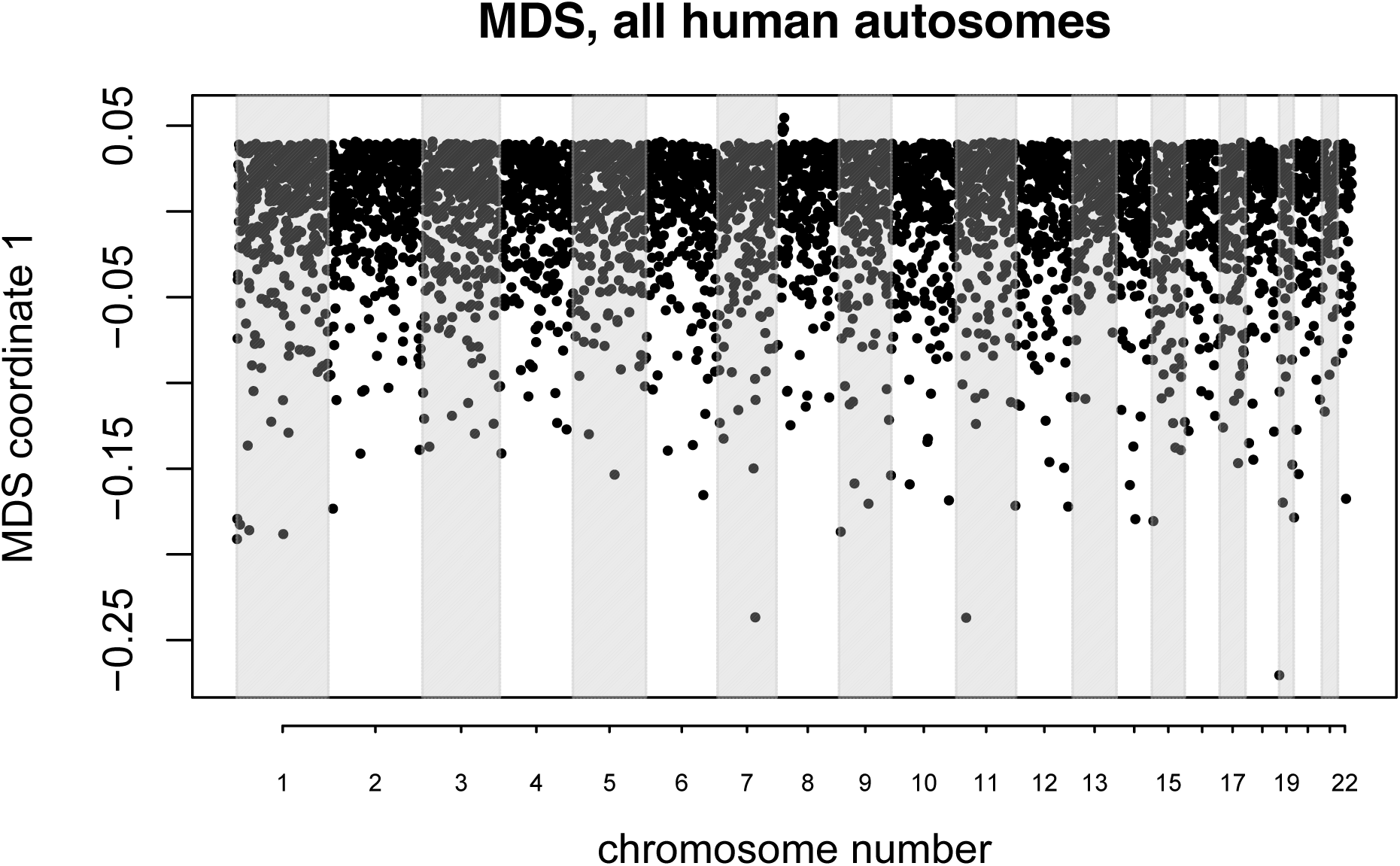
MDS visualization of variation in the effects of population structure amongst windows across *all* human autosomes simultaneously. The small group of windows with positive outlying MDS values lie around the inversion at 8p23.

**Figure S17:**
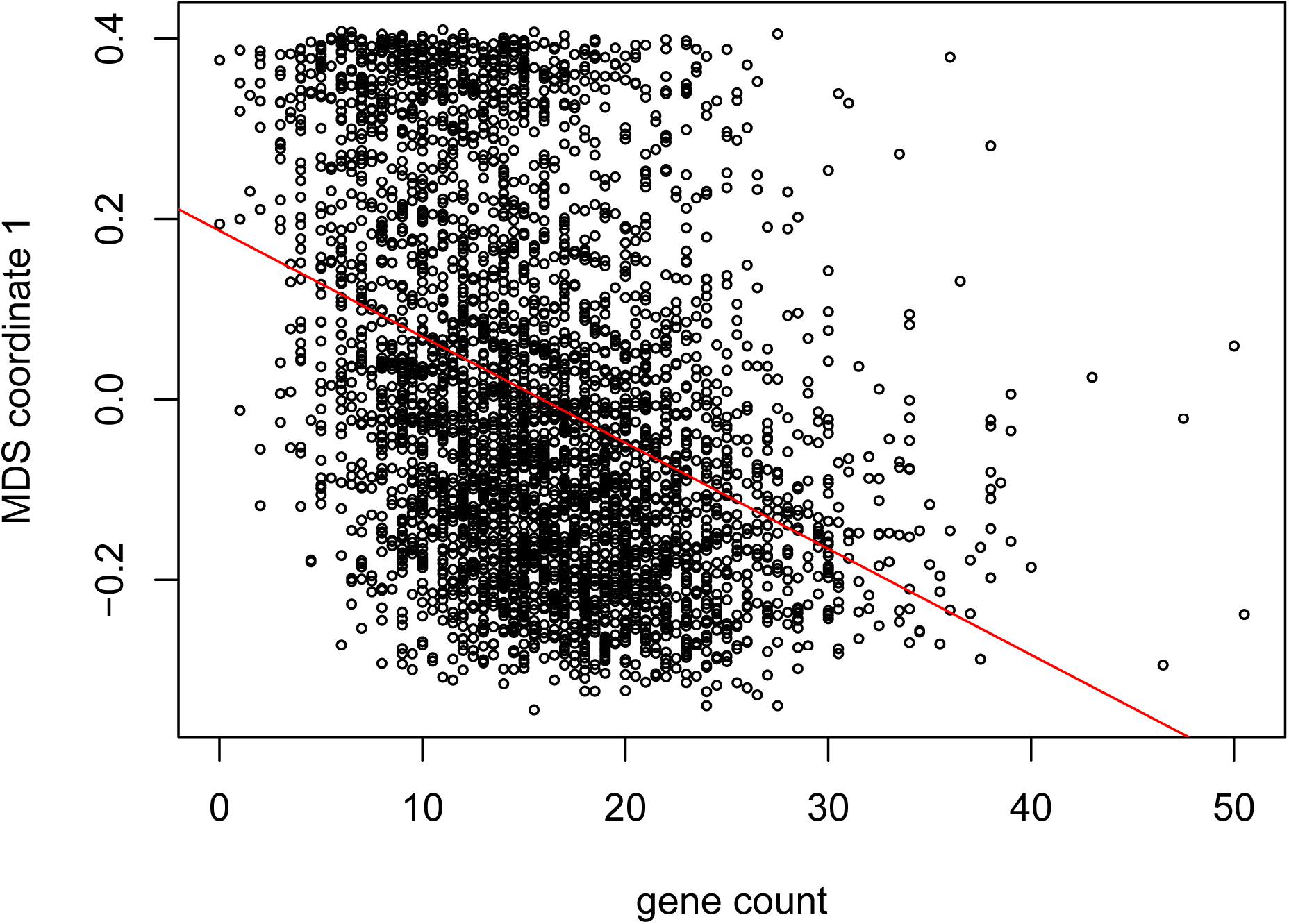
First MDS coordinate against gene density for all 8 chromosomes of *M*. *truncatula*. The first MDS coordinate is significantly correlated with gene count (*r* = 0.149, *p* = 2.2 × 10^−16^).

**Figure S18:**
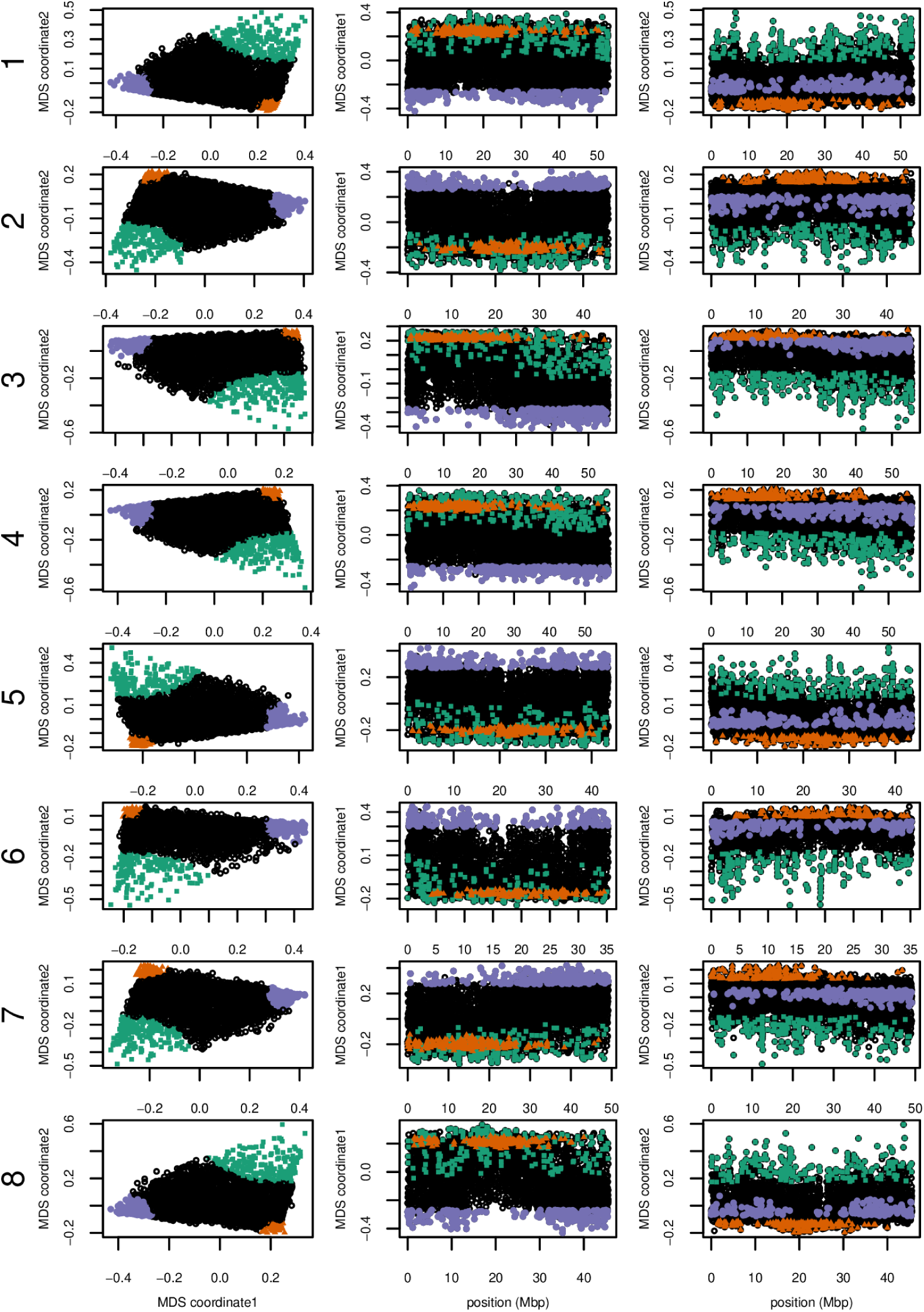
MDS visualizations of the effects of population structure for all 8 chromosomes of the *Medicago truncatula* data, using windows of 10^4^ SNPs.

**Figure S19:**
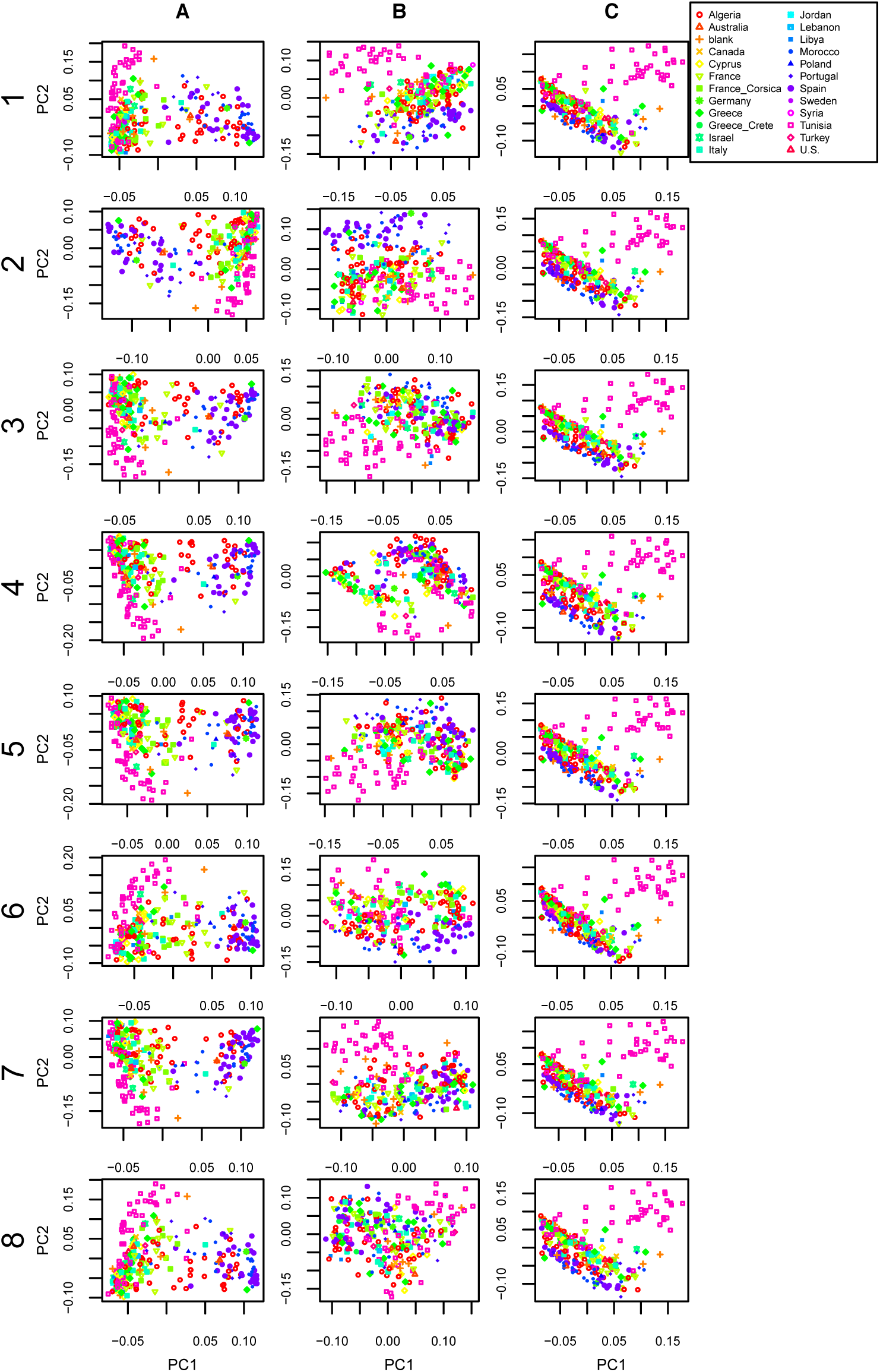
PCA plots for regions colored in Figure S18 on all 8 chromosomes of *Medicago truncatula*: (A) green, (B) orange, and (C) purple.

